# Form and function of actin impacts actin health and aging

**DOI:** 10.1101/2025.11.22.689949

**Authors:** Maxim Averbukh, Hunter Nelson, Tiffany Wang, Caitlin M Lange, Athena Alcala, Tripti Nair, Juri Kim, Daniella Suh, Matthew Vega, Shihong Max Gao, Naibedya Dutta, Rebecca Aviles Barahona, Aeowynn J Coakley, Joe A. Gerke, Sofia F. Odron, Jacqueline G. Clarke, Navjot Singh, Matthew S. Nickel, Jacob C. Peterson, Hannah L. Lam-Truong, Darius Moaddeli, Toni Castro Torres, Maria Oorloff, Melissa Peruch, Hemal Mehta, Max A. Thorwald, Sean P. Curran, Meng C Wang, Tyler A. Johnson, Eileen M. Crimmins, Thalida Em Arpawong, Caroline Kumsta, Gilberto Garcia, Ryo Higuchi-Sanabria

## Abstract

The actin cytoskeleton is a fundamental and highly conserved structure that functions in diverse cellular processes, yet its direct contribution to organismal aging remains unclear. Here, we systematically interrogated how genetic and pharmacologic perturbations of actin structure and function influence lifespan and various hallmarks of aging in *Caenorhabditis elegans*. Whole-animal and tissue-specific knockdown of actin and key actin-binding proteins (ABPs) - *arx-2* (Arp2/3), *unc-60* (cofilin), and *lev-11* (tropomyosin) - led to premature disruption of filament organization, reduced lifespan, and tissue-specific physiological defects. Bulk and single-nucleus RNA-sequencing revealed that ABP knockdowns elicited a strongly “aged” transcriptome. Actin dysfunction broadly exacerbated many age-associated phenotypes, including mitochondrial dysfunction, lipid dysregulation, loss of proteostasis, impaired autophagy, and intestinal barrier failure. Pharmacological destabilization with Latrunculin A mirrored genetic knockdowns, while mild stabilization with Jasplakinolide modestly extended lifespan, emphasizing that optimal and finely-tuned actin function is critical for healthy aging. Finally, analysis of human genome-wide association data revealed that common *ACTB* polymorphisms correlate with differences in age-related decline in gait speed, suggesting evolutionary conservation of actin’s role in healthy aging. Taken together, our results provide a comprehensive and publicly accessible resource that maps, for the first time, how actin integrity intersects with diverse aging pathways across tissues and scales. This descriptive framework is intended to enable future mechanistic discovery by offering a deep, unbiased dataset that can be integrated with emerging studies to define how actin dynamics contribute to aging.

## Introduction

The actin cytoskeleton is a dynamic three-dimensional scaffold of protein filaments and is fundamental to the structure, function, and dynamic adaptability of all eukaryotic cells^1^. Its evolutionary conservation demonstrates its critical role in many essential cellular processes in virtually all single- and multi-cellular eukaryotes, including cell division, motility, intracellular transport, and autophagy^2^. On a physiological level, actin’s integrity and dynamics are required for muscle contraction, neuronal plasticity, immune responses, tissue development, gut barrier integrity, and many other critical processes in multicellular organisms. Given its central role in these essential processes, the precise regulation of actin dynamics is critical for organismal health, and its dysregulation is found in many human pathologies, including in age-related diseases such as neurodegeneration, cancer, and muscle myopathies^2^. However, the contribution of the actin cytoskeleton in general aging pathology is only beginning to be understood.

Despite being synthesized by polymers of only a handful of proteins encoded by genes with >90% sequence identity (e.g., ACT-1 through ACT-5 in *C. elegans*; and α, β, and γ in mammals), the actin cytoskeleton can create diverse structures with vastly different functions within each cell type and tissue. Actin’s functional versatility is orchestrated by actin-binding proteins (ABPs) that coordinate assembly, disassembly, and organization of actin multi-protein complexes into higher-order structures capable of driving distinct biological processes^3^. Key among these regulators are the Arp2/3 complex that nucleates branched actin networks that drive cell protrusion^4^; cofilin, which severs filaments and promotes actin turnover^5^; formins which nucleate and elongate linear actin filaments^6^; and tropomyosin/troponin, which stabilizes filaments and is crucial for muscle function^7^. The interplay between these ABPs allows cells to tailor the form and dynamics of the actin cytoskeleton for diverse and specific functions, such as muscle contractions, tight junctions in the intestine, and wound healing. However, the mechanisms that preserve cytoskeletal integrity and function throughout an organism’s lifespan, particularly under stress and during aging, are not fully understood^2^.

Research in model organisms has provided significant insights into the relationship between actin homeostasis, aging, and longevity. In *C. elegans*, aging is associated with a progressive decline of actin filament stability^8^. Interventions that preserve actin integrity, such as overexpression of the genes encoding the heat shock transcription factor HSF-1 or the bromodomain protein BET-1, have been shown to extend lifespan and healthspan, providing a direct link between actin maintenance and organismal aging. HSF-1, for instance, can upregulate actin-regulatory genes like *pat-10*, contributing to its pro-longevity effects^9^. Similarly, BET-1 transcriptionally upregulates actin regulatory genes, thereby increasing actin stability and delaying age-associated disorganization of actin, contributing to lifespan extension^2^. In *Drosophila melanogaster*, one study showed that an age-associated accumulation of filamentous actin (F-actin) in the brain impairs autophagy, which leads to neuronal dysfunction and cognitive decline^10^. Reducing this F-actin accumulation by targeting the formin-like gene *Fhos* can restore autophagy and extend healthspan^10^.

In vertebrate systems, the consequences of actin dysfunction are most well understood in the context of neurodegeneration, muscle dysfunction, immune dysregulation, and cancer. The actin cytoskeleton is required for neuronal development, the morphogenesis of dendritic spines, and the synaptic plasticity underlying learning and memory^11–13^. Abnormal cofilin activity and the formation of cofilin-actin rods are implicated in neurodegenerative diseases like Alzheimer’s^14^. In the immune system, dynamic actin rearrangements drive immune cell migration, phagocytosis, and the formation of the immunological synapse necessary for T cell activation and cytotoxic responses^15,16^. Cardiovascular health also relies on actin integrity for endothelial barrier function and vascular smooth muscle contractility, with regulators like profilin-1 implicated in hypertension and atherosclerosis^17–21^. Moreover, the actin cytoskeleton is involved in cancer progression, driving cell motility, invasion, and immune evasion^22–24^. Importantly, aging is a major risk factor for all these actin-related diseases, providing further evidence of the association between actin and aging.

Although many studies to date correlate and associate alterations of actin to effects on aging, there is a lack of direct studies that display targeted disruption or protection of actin and their impact on aging. Therefore, to measure the impact of actin dysfunction on the aging process we targeted the actin cytoskeleton using various genetic and chemical methods in *C. elegans*. First, we induced actin dysfunction using whole animal and tissue-specific RNA interference (RNAi): 1) non-lethal knockdown of the actin gene itself (10% *act-1* RNAi, which targets all five isoforms of actin and does not cause developmental arrest as previously described^2^) for global actin dysfunction, and more targeted approaches of 2) *arx-2* knockdown to target Arp2/3 and inhibit growth of branched actin networks 3) *lev-11* knockdown to target tropomyosin and destabilize actin filaments and 4) *unc-60* knockdown to target cofilin and inhibit severing of actin filaments and actin seeding. To complement these genetic manipulations, we used two global actin-targeting marine-derived chemical probes: 1) latrunculin A (LatA) – that severs and depolymerizes actin filaments, while also sequestering available actin monomers and 2) jasplakinolide (Jasp) – which promotes actin filament polymerization and stabilization.

This work is intended as a foundational resource rather than providing mechanistic models: we performed an expansive and unbiased study analyzing how altering actin in various ways impact numerous metrics of cellular, transcriptomic, and physiological health during aging, including studying tissue-specific effects of actin alterations on aging.

## Results

### Systemic RNAi of actin regulatory genes causes premature aging phenotypes

Several studies have shown that actin cytoskeletal organization and integrity declines during normal aging^2,8,9^. To determine whether changes to actin are a causative driver of aging or just a consequence of aging, we sought to determine the impact of actin dysfunction on metrics of aging. We first utilized RNAi to knockdown genes encoding actin (*act-1*, which targets all 5 isoforms of actin^2^, and will be referred to simply as “actin knockdown” for simplicity) and ABPs (*arx-2, lev-11, unc-60*). To confirm that these genetic perturbations lead to an accelerated disruption of actin filaments during aging, we used LifeAct::mRuby-expressing animals to visualize actin structure and organization in young, mid-age, and old animals in a tissue-specific manner. LifeAct is a 17-amino-acid fragment of a yeast actin-binding protein, Abp140p, which binds directly to actin filaments and is often used to visualize actin filaments in live cells/organisms by fusing it to a fluorescent protein^25^. We utilized previously-developed stable transgenic lines expressing LifeAct::mRuby in various tissues^2^.

Actin filaments in muscle cells are organized in linear striations parallel to muscle fibers, which become disorganized at approximately day 9 of adulthood, and display fragmentation and severe disruption at late age (∼day 13-15). In the intestine, actin is visualized as dense cables that wrap around intestinal cells to make up the gut barrier, which begin to display disorganization and mislocalization at mid-age at approximately day 5 of adulthood. In the hypodermis, actin filaments are sparsely localized throughout the hypodermis at day 1 and become organized into star-like structures that resemble endocytic vesicles coated with actin around day 3-4, which reduce in number and are eventually completely lost during the aging process^8^. Upon RNAi knockdown of actin and ABPs, we saw premature onset of aging phenotypes at much earlier time-points (**Fig. 1**). Interestingly, we saw that RNAi knockdown of ABPs had variable effects in each tissue. For example, *unc-60* knockdown resulted in mislocalization and disorganization of actin in the intestine even at day 1 of adulthood, whereas *arx-2* and *lev-11* knockdown displayed similar timing in actin disorganization (day 5 to day 9) compared to wild-type, but displayed much stronger phenotypes with age (**Fig. 1A-B**). In contrast, actin knockdown resulted in almost no observable actin filaments in the intestine at day 9. Similarly, actin knockdown resulted in no observable actin filaments in the hypodermis, and *arx-2* knockdown phenocopied these effects. However, *lev-11* and *unc-60* knockdown resulted in only a minor reduction in the observable amount of actin structures in the hypodermis (**Fig. 1C-D**). In the muscle, *act-1*, *arx-2*, and *lev-11* resulted in premature disorganization of actin structures, while *unc-60* had only mild effects (**Fig. 1E-F**). Altogether, these data show that RNAi knockdown of actin or ABPs reliably induce premature disruption of actin structures during aging, though at variable levels in each tissue.

**Fig 1:**
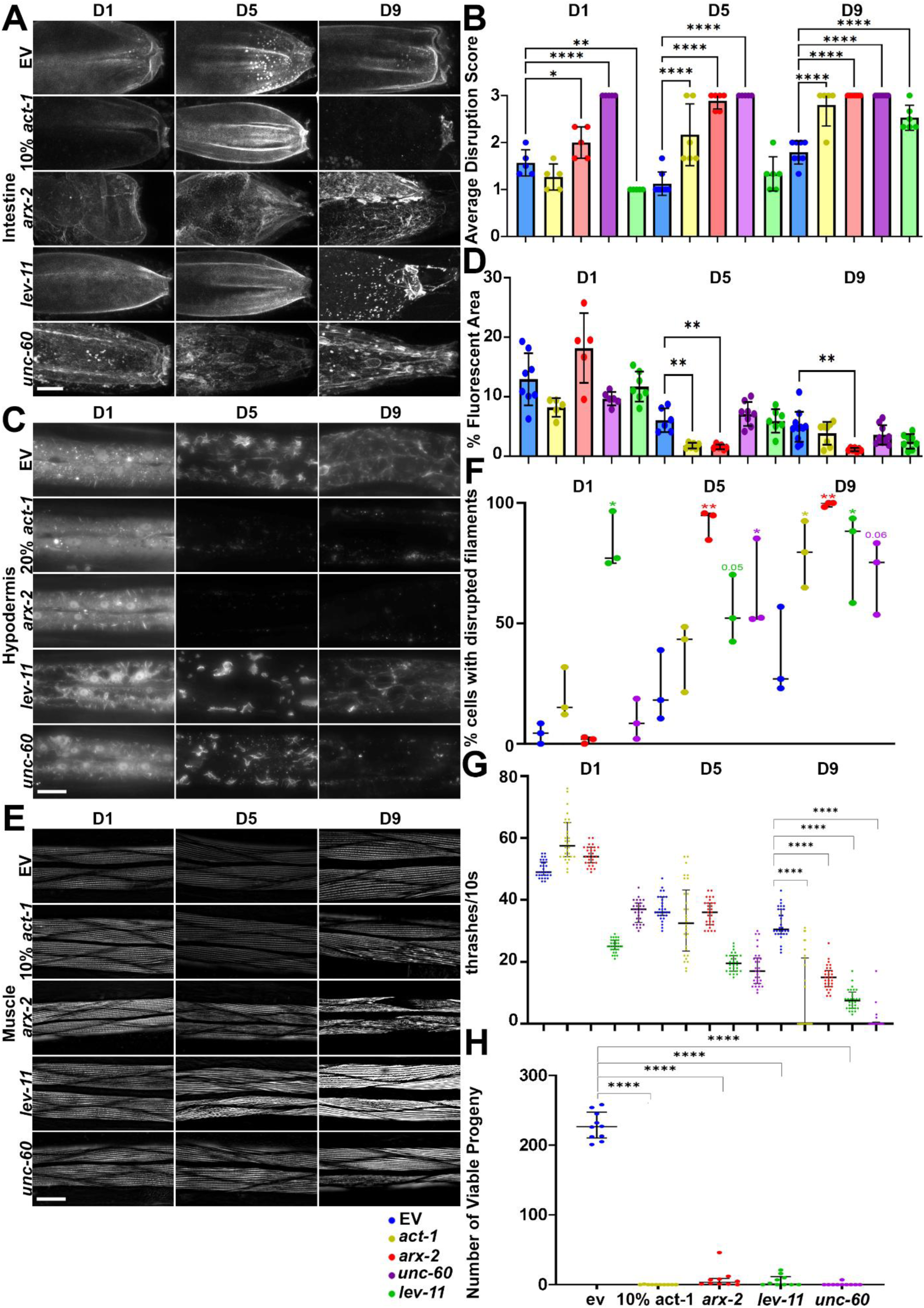
Knockdown of ABPs decreases lifespan and leads to accelerated actin disruption. **(A-E)** Representative fluorescent images of adult animals expressing LifeAct::mRuby from an intestine-specific (*gly-19p*) **(A)**, hypodermis-specific (*col-19p*) **(C)**, or muscle specific (*myo-3p*) promoter **(E)** and quantification of **(B)** intestinal, **(D)** hypodermal and **(F)** muscle actin filament disruption. Animals were grown on empty vector (EV), a 1:9 or 2:8 mix of *act-1*:EV (10% *act-1* or 20% *act-1*), *arx-2*, *lev-11*, or *unc-60* RNAi from hatch. All animals were imaged on day 1, 5, and 9 of adulthood and images were captured on a Leica Stellaris (intestine) or Leica Thunder Imager (hypodermis, muscle). Scale bar is 10 µm. **(G)** Thrashing assays were performed on N2 animals grown on EV (blue), a 10:90 mix of *act-1*:EV (10% *act-1*, yellow), *arx-2* (red), *lev-11* (green), and *unc-60* (purple) RNAi from hatch on day 1, 5, and 9 of adulthood. Recordings were performed in M9 solution and thrashing was manually scored over a 10 second recording. Data is representative of three independent trials. N = 30-33 worms per condition. Each dot represents a single animal, and lines represent median and interquartile range. *** = p < 0.001 calculated using non-parametric Mann-Whitney testing. **(H)** Brood assays were performed on wildtype N2 animals grown on empty vector (EV, blue), a 10:90 ratio of *act-1*/EV (10% *act-1*, yellow), *arx-2* (red), *lev-11* (green), or *unc-60* (purple) RNAi from hatch. Brood assays were measured as any viable animal that was hatched from eggs laid within 48 hours.. **** = p < 0.0001 calculated using non-parametric Mann-Whitney testing.

To determine the physiological and functional consequences of actin, we measured the effects of actin and ABP knockdown on lifespan and healthspan metrics. Since actin in the muscle is required for muscle contractions and animal motility^26^, we performed a liquid thrashing assay to measure maximal animal motility during aging^27^. We found that knockdown of actin or ABPs result in a significant decrease in locomotor behavior at older ages (**Fig. 1G**), suggesting that muscle function is significantly perturbed in all conditions. Strikingly, knockdown of actin and *arx-2* resulted in an increase in motility at day 1, suggesting a potential compensatory mechanism is activated in early adulthood, which collapses during aging. We found that perturbations of actin resulted in severely reduced brood size (**Fig. 1H**), which is consistent with actin function being required for proper cell division both during germline formation and throughout animal development^28–32^. Finally, RNAi knockdown of actin or any ABP led to a significant decline in lifespan (**Fig. 2A-B**).

**Fig 2:**
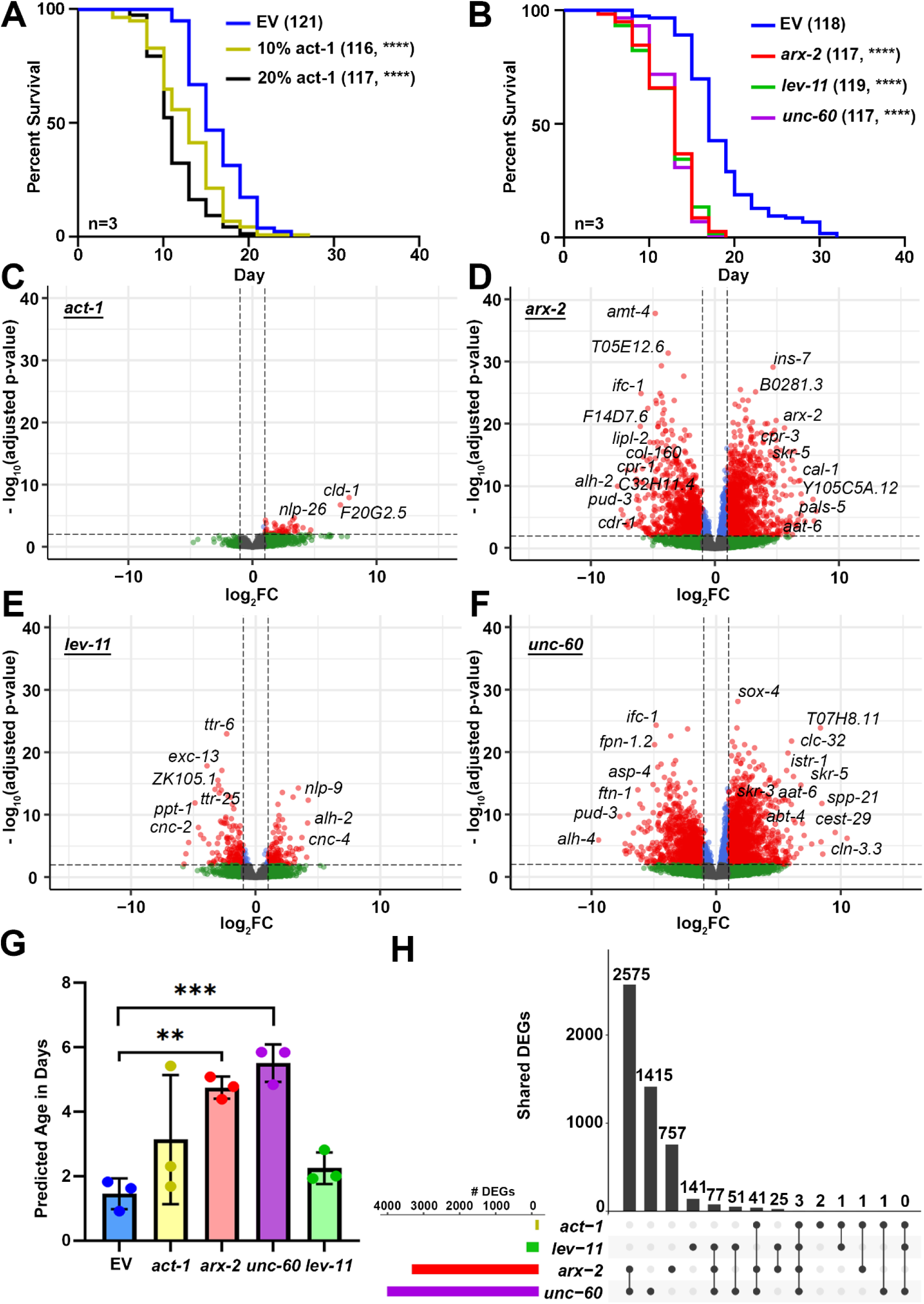
Whole organism RNAi of actin regulatory genes induces an aged transcriptome. **A)** Lifespans of wild-type N2 animals grown on empty vector (EV, blue) or varying ratios of *act-1*:EV RNAi: 1:9 (yellow), or 2:8 (black) from hatch. **(B)** Lifespans of wild-type N2 animals grown on empty vector (EV, blue), *arx-2* (red), *lev-11* (green), or *unc-60* (purple) RNAi from hatch. See **Table S1** for lifespan statistics. Volcano plots of genome-wide changes in gene expression upon RNAi knockdown of **(C)** *act-1,* **(D)** *arx-2*, **(E)** *lev-11* and **(F)** *unc-60*. Red dots indicate significantly differentially expressed genes with p-value ≤ 0.01 and log_2_FC>|2|. Blue dots indicate significantly differentially expressed genes with p-value ≤ 0.01 and log_2_FC<|2|. Green dots indicate significantly differentially expressed genes with p-value ≥0.01. See **Table S3** for a list of differentially expressed genes and expression values. **(G)** Predicted Age of day 1 adult worms, using BiT Age algorithm. ** = p < 0.01, *** = p < 0.001 calculated using paired t-test. . See **Table S4** for the BiTAge output. **(H)** An UpSet plot showing the number of all differentially expressed genes (p-value ≤ 0.01) unique to and shared between all tested conditions.

### Whole organism RNAi of actin regulatory genes induces an aged transcriptome

The actin cytoskeleton plays an important role in several processes that affect gene expression including chromatin remodeling and transcription^33,34^. Therefore, to determine whether actin dysfunction drives global changes to gene expression, we first performed transcriptomics analysis using whole-animal, bulk RNA-seq under RNAi of actin and ABPs of Day 1 adult animals. RNA-seq libraries showed Phred scores >35 with consistent mapping and clustering across replicates, indicating high-quality data (**Fig. S1A-D**). Knockdown of *arx-2* and *unc-60* led to sizable changes in gene expression, while much milder changes were seen in actin and *lev-11* knockdown (**Fig. 2C-F**). It is possible that this is due to the ubiquitous expression – and therefore general requirement – of *arx-2* and *unc-60* throughout the entire organism. In contrast, *lev-11* is primarily expressed in the muscle, thus likely having a smaller impact on bulk transcriptome. Finally, actin knockdown is lethal and thus needs to be titrated to a small dosage (10-20% knockdowns used in this study), and thus likely has a much milder effect. Consistent with actin dysfunction driving premature aging, we also observed acceleration of transcriptomic age, measured using the BiT age algorithm^35^, upon knockdown of *arx-2* and *unc-60*, which was not seen in actin or *lev-11* knockdowns (**Fig. 2G**). To determine whether knockdown of a single actin regulatory gene results in a compensatory upregulation of other actin regulatory genes, we next profiled gene expression changes of all major actin regulatory genes. To our surprise, most actin regulatory genes were either downregulated or unaffected in each of our knockdown conditions (**Fig. S1E**), (note: although *arx-2* appears to be upregulated in the *arx-2* knockdown condition, this is likely an artifact of the RNAi construct being sequenced).

Next, we performed an upset plot analysis to determine whether a common transcriptomic signature exists in response to actin dysfunction. To our surprise, most knockdown conditions exhibited highly variable changes to overall transcriptomes, with only *arx-2* and *unc-60* knockdown showing any significant overlap (**Fig. 2H**). The comparison between the differentially expressed genes (DEGs) of *arx-2* and *unc-60* knockdown conditions showed very high concordance between the groups, with almost all DEGs being shared (**Fig. 3A-B**). Although *arx-2* and *unc-60* also show significant concordance with *lev-11*, the number of genes overlapping between these groups are relatively small (<200) compared to the overlap between *arx-2* and *unc-60* (>1500), which is consistent with these two genes being ubiquitously expressed and likely being required for all tissues, whereas *lev-11* is a muscle-enriched tropomyosin (**Fig. 3C-F**). Further Gene Ontology (GO) analysis revealed that a shared general response to actin disruption is clear changes related to immune and defense response (**Fig. S2**), highlighting that at least one signature of genes is shared across all conditions. Consistent with this hypothesis, GO analysis of overlapping upregulated DEGs between *arx-2* and *unc-60* are primarily immune-response related genes (**Fig. 3G**). In contrast, GO analysis of commonly downregulated DEGs between *arx-2* and *unc-60* revealed primarily change to metabolic processes (**Fig. 3H**). Since *lev-11* knockdown displayed the most unique gene expression changes, we performed GO analysis of unique DEGs and found, as expected, that a proportion of gene expression changes are related to muscle function (**Fig. 3I**). Interestingly, neuropeptide signaling was the most enriched GO term, although this is consistent with a role for neuronal signaling across neuromuscular junctions to coordinate proper muscle contraction^36,37^.

**Fig 3:**
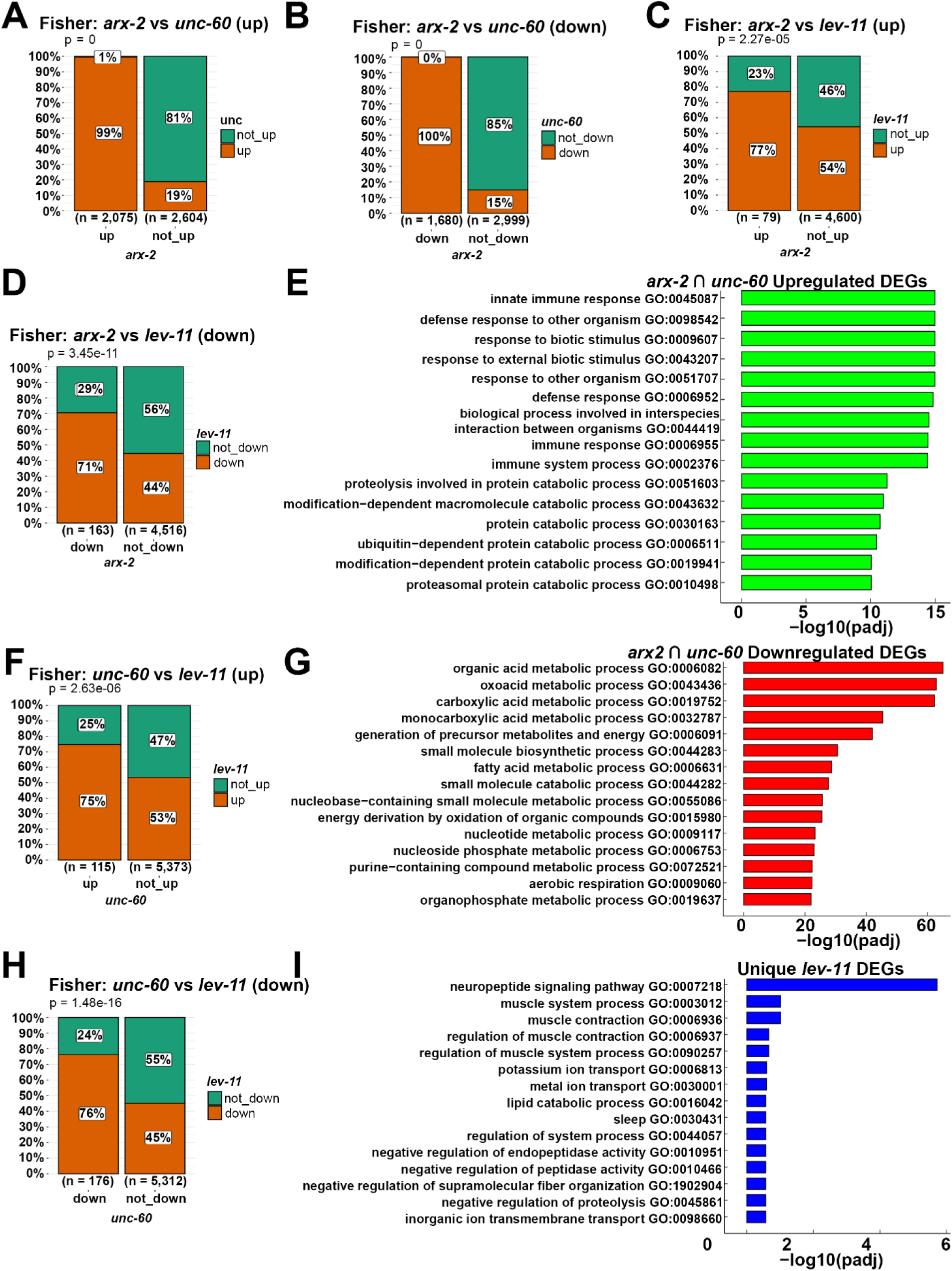
Knockdown of various ABP results in highly concordant transcriptomic changes. Bar plots to visualize the overlap of up- and down-regulated differentially expressed genes (DEGs) between animals treated with **(A)(B)** *arx-2* and *unc-60*, **(C)(D)** *arx-2* and *lev-11*, and **(E)(F)** *unc-60* and *lev-11* using ggbarstats function in R. Statistical significance of the overlap was assessed using Fisher’s exact test. GO analysis of DEGs **(G)** upregulated and **(H)** downregulated exclusively in both *arx-2* and *unc-60* conditions. **(I)** GO analysis of DEGs unique to the *lev-11* condition. See **Table S5** for Go IDs, genes and statistics of the GO Analysis.

### Systemic RNAi of actin regulatory genes exacerbates biological hallmarks of aging

Since we saw changes to whole animal health and longevity upon disruption of the cytoskeleton, we next sought to determine the impact of actin disruption on cellular metrics of health and aging. Several biological hallmarks of aging^38^ are routinely studied in *C. elegans*, including mitochondrial dysfunction, nutrient dysregulation, dysbiosis, proteostasis, and autophagy. In addition, the actin cytoskeleton has been shown to be involved in regulation of many of these pathways, although the direct relevance of actin to these cellular mechanisms and their impact on aging have not been extensively studied, with few exceptions. Therefore, we directly measure how actin dysfunction impacts many age-associated cellular processes.

Mitochondrial function deteriorates during aging in almost all organisms, including *C. elegans*, mice, and humans^38–41^. Actin plays crucial roles in regulation of mitochondrial health, including in mitochondrial fission, fusion, motility, and mitochondrial damage response^42–50^. Therefore, we visualized effects of actin and ABP knockdown on mitochondrial morphology, using a matrix–localized green fluorescent protein (MLS::GFP) in the intestine, hypodermis, and muscle^51^. Actin disruption resulted in dramatic changes in mitochondrial morphology in the intestine, muscle, and hypodermis (**Fig. S3A-F**). Our qualitative mitochondrial images show that knockdown of actin, *arx-2*, and *unc-60* resulted in increased fragmentation of mitochondria during aging, while *lev-11* knockdown resulted in a more hyperfused network.

To quantify mitochondrial morphology, we utilized mitoMAPR, which provides automated quantification of mitochondrial length, junction points, object number, and mitochondrial footprint (total number of objects detected and total area occupied by mitochondria within a defined region)^52^. While mitoMAPR can robustly quantify mitochondrial length in an unbiased manner, there are some limitations with the skeletonization of images required for quantification, particularly when major changes to fusion and fission occur^53,54^. Therefore, our qualitative data did not always match quantitative analyses with mitoMAPR, except in some conditions (**Fig. S3B, D, F**).

Therefore, as a more direct measure to determine whether our observed qualitative changes in morphology reflected quantifiable, functional changes in mitochondria, we measured mitochondrial respiration rates in whole animals using a previously validated Seahorse assay^55^. Consistent with changes to mitochondrial morphology, disruption of actin resulted in a significant reduction in oxygen consumption rate (OCR), suggesting that the changes in mitochondrial morphology had direct impact on mitochondrial function (**Fig. S3G**). Interestingly, *lev-11* knockdown had the most profound effect on OCR. However, since *lev-11* knockdown results in a significant defect in muscle function even at day 1 (**Fig. 1G**), it is possible that *lev-11* knockdown also disrupts non-mitochondrial respiration, as this is primarily driven by muscle function in *C. elegans*^55–57^, and *lev-11* knockdown animals exhibit severely dysfunctional muscle function. Indeed, we saw that even with sodium azide treatment that eliminates mitochondrial respiration, *lev-11* knockdown still reduced OCR, suggesting that *lev-11* knockdown reduces both mitochondrial and non-mitochondrial respiration.

Given that mitochondria are central hubs for lipid synthesis and β-oxidation, and their disruption can impair lipid homeostasis, we next hypothesized that cytoskeletal defects may indirectly contribute to lipid dysregulation. Therefore, we used a lipid droplet marker DHS-3::GFP to measure lipid droplet levels in the intestine under actin disruption throughout the lifespan^58,59^.

Interestingly, knockdown of the ABPs *arx-2*, *unc-60* and *lev-11* all resulted in a decrease in lipid droplet levels at mid and old age (**Fig. S4A-B**). While all conditions show a decreasing trend in DHS-3::GFP signal, *unc-60* and *lev-11* show the most profound statistically significant decrease in signal. However, since DHS-3::GFP can exist both within lipid droplets and within the cytoplasm, we completed high-resolution confocal microscopy of the DHS-3::GFP marker to investigate specific changes in lipid droplet number, size, and morphology **(Fig. S4C)**.

Interestingly, there is not a distinct decrease in the number of lipid droplets present in animals with ABP knockdown, but trend towards an increase, with only day 1 animals hitting statistical significance **(Fig. S4D)**. However, the size of lipid droplets present is drastically decreased throughout the aging process under actin disruption **(Fig. S4E)**. Low magnification imaging showed a decrease in signal likely due to the decrease in the size of lipid droplets. Altogether, our data suggest that actin and ABP knockdown results in dysregulation of lipid homeostasis, which is further supported by our RNAseq analysis, where we found differential expression of many lipid regulatory genes (**Fig. S4F**).

Another central cellular function that declines with age but is necessary for organismal health is protein homeostasis, or proteostasis. Specifically, degradation of damaged or misfolded proteins is key to maintaining proteostasis. Actin has been recently implicated as a key player in proteostasis maintenance, specifically through proteasome regulation^60^. Importantly, actin disruption can exacerbate aggregation of proteins in neurons to drive neurodegenerative pathology. Therefore, we visualized the human aggregation-prone protein, polyQ40 fused to yellow fluorescent protein expressed specifically in neurons^61^. Here, we used the pan-neuronal Q40 strain, which has a very clear phenotype under neurotoxic conditions where neuronal aggregate-like foci are jettisoned into the body cavity^62^. Perturbing actin by knocking down *unc-60* had the most profound, statistically significant increase in the number and size of foci throughout the aging process (**Fig. S5A-C**). Interestingly, *lev-11* knockdown had a very similar increase in number and size of neuronal foci, despite *lev-11* primarily expressed in the muscle (**Fig. S5A-C**), suggesting that either disruption of muscle health can indirectly impact neurons, or there exist a muscle to neuron signaling axis that can affect protein homeostasis pathways.

This is consistent with our transcriptomics data showing that *lev-11* knockdown resulted in differential expression of neuropeptide related genes (**Fig. 3I**). Interestingly, *arx-2* knockdown led to a significant increase of aggregation at day 1 of adulthood but showed no increase in aggregates at later age (**Fig. S5A-C**). However, due to the severely shortened lifespan of *arx-2* knockdown animals, it is possible that survivorship bias may confound these late-life results. Notably, the pan-neuronal Q40 strain is a systemic RNAi background in which neurons are largely resistant to RNAi, suggesting that the observed effects on proteostasis are likely cell-nonautonomous.

One major pathway for clearance of protein aggregates is through autophagy. In addition, dynamic remodeling of actin is essential for multiple stages of the autophagy process, including autophagosome formation, cargo sequestration, and vesicle trafficking^63,64^. Actin filaments not only provide the mechanical force and scaffolding necessary for membrane deformation but also coordinate the spatial organization of autophagic machinery^65,66^. Therefore, to investigate the impact of actin dysfunction on autophagy, we first used *C. elegans* expressing a GFP-tagged LGG-1/Atg8 reporter to visualize autophagosomes^67^. We found that actin disruption led to significant changes in the quantity of autophagosomes in a tissue specific manner. In the intestine actin*, unc-60* and *arx-2* knockdown all led to a significant increase in the number of LGG-1 puncta, while *lev-11* knockdown led to a surprising decrease in puncta (**Fig. S6A-B**).

Interestingly, we saw the opposite results in the muscle cells where actin*, unc-60* and *arx-2* knockdown all led to a significant decrease in the number of LGG-1 puncta, while *lev-11* knockdown had no significant effect (**Fig. S6B,D**). Surprisingly, none of the knockdown conditions had any effect on LGG-1 puncta in hypodermal seam cells (**Fig. S6B-C**). We further checked changes in expression of autophagy genes in our bulk sequencing dataset and found that knockdown of *arx-2* and *unc-60* led to an upregulation, while knockdown of *act-1* and *lev-11* led to a decrease in expression (**Fig. S6E**). These data suggest that similar to changes in actin organization, RNAi knockdown of different actin regulatory components can have differential effects on autophagy in each tissue.

Lastly, one of the core causes and predictors of mortality across species is intestinal barrier dysfunction^68^. In *C. elegans*, the intestinal lining contains actin-mediated tight junctions, which breakdown during aging leading to bacterial colonization of the intestine^69^. Our previous work has established that stabilizing actin filaments by *bet-1* overexpression protects *C. elegans* from bacterial colonization at later age^2^. Moreover, our bulk RNA-seq revealed that there is a large transcriptomic response of immunity-related genes in all our knockdown conditions (**Fig. S2**). Therefore, we first confirmed that direct perturbations of actin impact gut barrier integrity. Utilizing a well-established fluorescent bacteria colonization assay, we found that RNAi knockdown of actin or ABPs resulted in a significant increase in gut colonization at day 1 of adulthood compared to an empty vector control (**Fig. S7A-B**). However, these differences diminished with age, and only *arx-2* knockdown showed an increased colonization at day 9 of adulthood, likely due to control animals also exhibiting defects in gut barrier integrity at old age^69^ and thus becoming indistinguishable from actin dysfunction conditions.

### Cell-type specific effects of actin dysfunction

We found that RNAi knockdown of ABPs had variable effects in each tissue tested (**Fig. 1**), which raised an intriguing hypothesis that each individual cell type has a different requirement for actin regulatory proteins. This is unsurprising considering the vastly different structures and function of actin in each tissue. To further test the cell-type specific requirements of actin and ABPs, we utilized cell-type specific RNAi strains. These strains utilize a mutation in *rde-1*, the gene encoding the Argonaut machinery required to process RNAi, followed by cell-type specific *rde-1* rescue^70^. For neuron-specificity, we utilized a *C. elegans* strain with a mutation in the *sid-1* gene, encoding the double-stranded RNA importer also required for RNAi function, followed by neuron-specific *sid-1* rescue^71^. While whole-animal RNAi knockdown of actin is lethal and thus requires dilution of the RNAi to 10%^2^, cell-type specific actin knockdown was not lethal (**Fig. S7C**) and thus we tested both 10% and 100% actin RNAi for our cell-type specific studies. As expected, actin and ABP knockdown had variable effects when performed in each specific cell type (**Fig. 4**).

**Fig 4:**
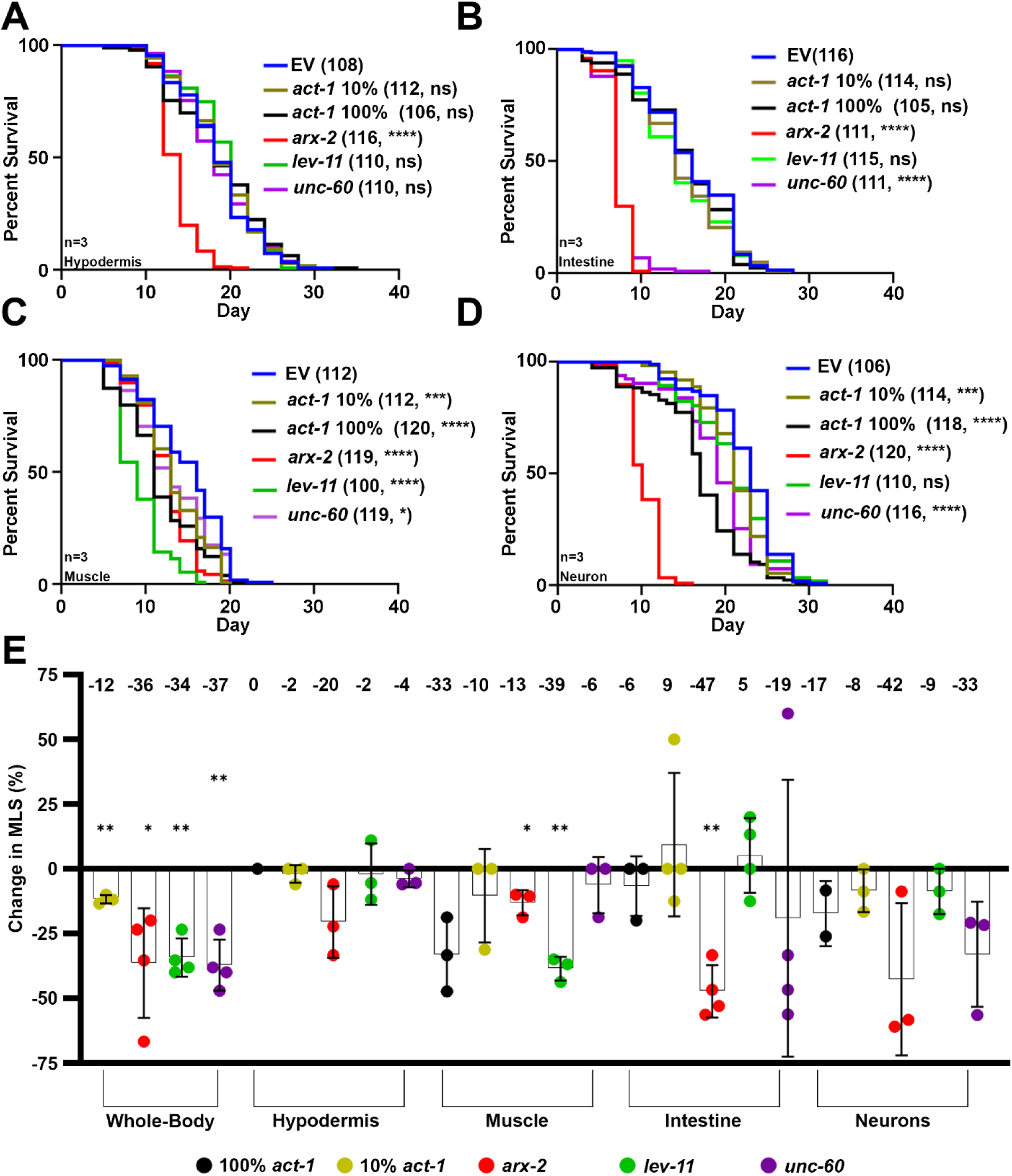
Actin disruption presents tissue-specific effects on lifespan. Lifespans of animals with functional RNAi only in the **(A)** hypodermis **(B)** intestine **(C)** muscle and **(D)** neurons grown on empty vector (EV, blue), 100% *act-1* (black), 10% *act-1* (yellow), *arx-2* (red), *lev-11* (green), or *unc-60* (purple) RNAi from hatch. **(E)** Average mean lifespan change (% MLS change) in wild-type animals and animals with tissue-specific functional RNAi after growth on RNAi of the listed ABPs compared to control. Horizontal values at the top represent the average percent change in median lifespan for each condition. Error bars indicate the s.d. * = p < 0.05, ** = p < 0.01 by a two-sided, one-sample t-test compared to hypothetical mean of 0. See **Table S1** for lifespan statistics.

For example, *lev-11* knockdown had the most dramatic effect in reducing lifespan in the muscle, with a much milder effect on the intestine and no effect in the hypodermis or neurons. This is consistent with *lev-11* being a muscle-enriched gene and its knockdown having the most dramatic effect on muscle actin morphology (**Fig. 1E-F**). *unc-60* knockdown had the most dramatic effect in reducing lifespan in the intestine, also consistent with our imaging data, which showed that *unc-60* knockdown resulted in disorganization of actin in the intestine as early as day 1 of adulthood (**Fig. 1A-B**). *unc-60* knockdown also exhibited a mild decrease in lifespan in the muscle and neuron but had very little effect in the hypodermis. Finally, *arx-2* knockdown was the only knockdown that consistently reduced lifespan in all tissues.

To further dissect the genetic mechanisms underlying cell-type specific effects of ABP knockdowns on health and longevity, we conducted single-nuclei RNA-sequencing (snRNA-seq) of systemic ABP knockdowns using a previously validated protocol^72^, followed by deconvolution of our bulk RNA-seq dataset^73^. Our snRNA-seq was able to collect thousands of nuclei for the largest tissues of the animals in most conditions **(Fig. S8A)** and showed robust cell clustering for each condition **(Fig. S8B-E)**. Although previous studies have shown that technical variability across *C. elegans* snRNA-seq datasets is minimal due to the contribution of hundreds of animals to data collection in a single sequencing run^72^, to ensure that our data were not skewed due to conducting only a single replicate, we utilized our snRNA-seq dataset to perform deconvolution of our bulk RNA-sequencing dataset. To first gain a holistic, zoomed-out view of the tissue-specific effects of systemic ABP knockdowns, we compared the transcriptional responses across tissues and targets. Spearman correlation analysis revealed some unexpected similarities and differences. Notably, muscle *lev-11* and *arx-2* knockdown and hypodermal *unc-60* and *arx-2* knockdown displayed a surprisingly high degree of similarity, suggesting that these tissues may share unexpected common mechanisms for maintaining cytoskeletal integrity or responding to actin dysfunction. In contrast, *lev-11* knockdown produced transcriptional profiles in the intestine that were distinct from those across all other tissues (**Fig. S9A)**. Interestingly, we found that the intestine was responsible for the largest transcriptional signature for each condition, potentially because it is more sensitive to RNAi and/or more reactive to actin disruption^74^(**Fig. S9B)**.

Next to determine whether cell-type specific phenotypic measures we identified in this study correlated with their tissue-specific changes in transcriptional profile, we investigated changes in gene expression of immune response, protein translation, and autophagy genes. We found that genes involved in immune response were differentially expressed in all tissues, with intestine showing the largest changes and muscle the least **(Fig. 5A)**. This was unsurprising considering that changes in immune response genes were one of the only shared responses across all conditions in our bulk datasets and all actin and ABP knockdown conditions affect gut barrier integrity. It has been reported that *C. elegans* exposed to stress can downregulate translation in response, however we did not find major changes in protein translation related genes in the intestine or in any other cell type^75–77^ (**Fig. 5B**). We also see comparable changes in expression of autophagy-related genes across tissue (**Fig. 5C**), suggesting that effects of actin disruption on autophagy may not be driven by transcriptional changes, but rather direct effects of altering actin dynamics on autophagy.

**Fig 5:**
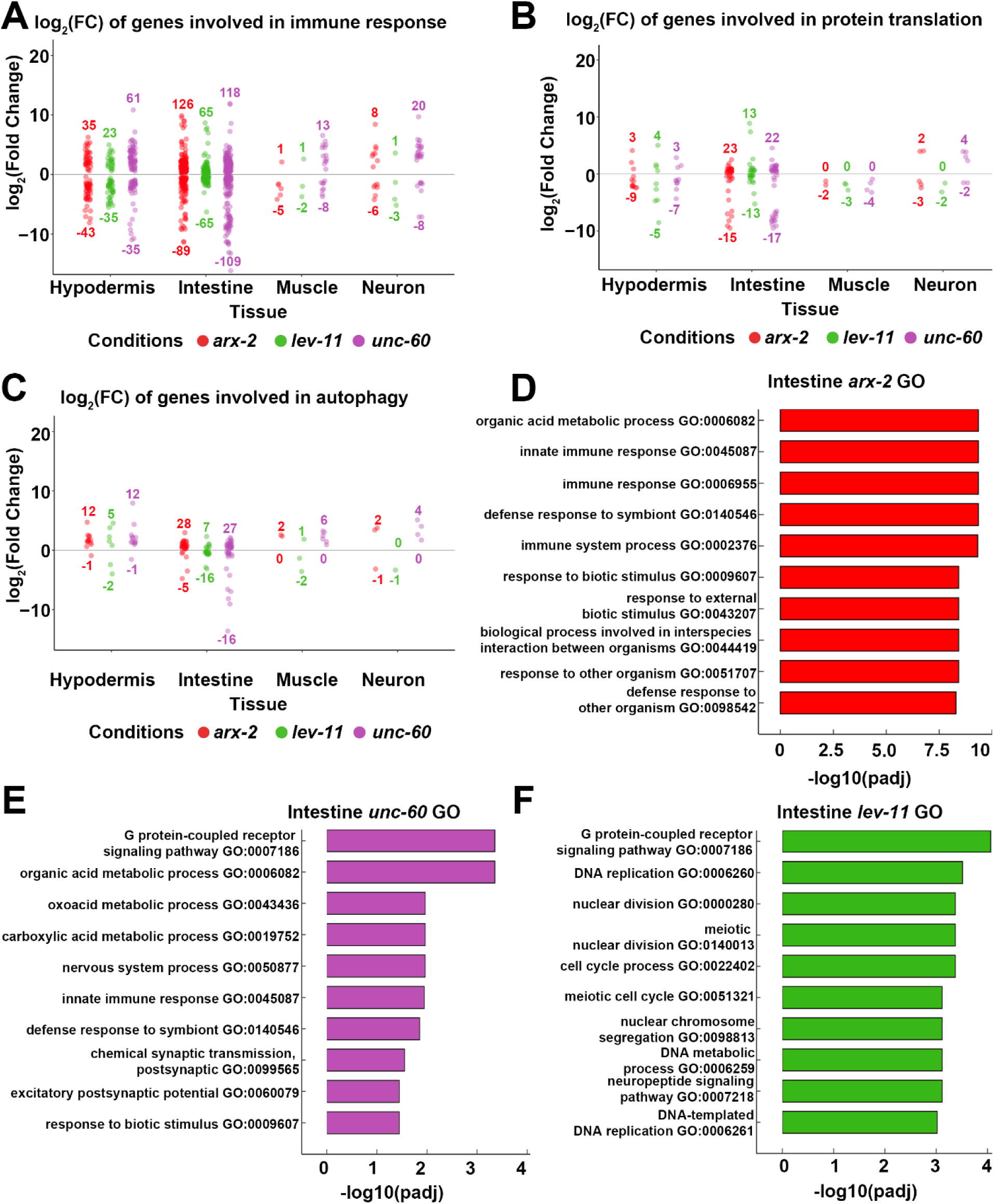
RNAi of ABPs induced cell-type specific transcriptional responses. **(A-C)** Dot plots showing significant DEGs involved in **(A)** immune response (annotated as involved in “immune response” in AmiGO2^188^), **(B)** protein translation (directly annotated as “regulation of translation” in AmiGO2^188^) and **(C)** autophagy (directly annotated as “autophagy” in AmiGO2^188^) under ABP knockdown in 4 key tissues: Hypodermis, Intestine, Muscle and Neuron. See **Table S6** for DEGs in each dotplot. **(D-F)** GO analysis of DEGs in the intestine under ABP knockdown. See **Table S7** for Go IDs, genes and statistics of the GO Analysis.

Similarly, we checked genes involved in other longevity-associated phenotypes tested in this study, including mitochondrial organization, lipid regulation, and general actin-related gene expression. As expected, many genes related to mitochondrial organization are downregulated in the hypodermis, intestine, and muscle upon ABP knockdown (**Fig. S9C**) which is consistent with mitochondrial fragmentation and loss of function in these conditions (**Fig. S3**). Interestingly, the hypodermis displays a similar number of significantly up- and downregulated genes under ABP knockdown, whereas neurons display primarily an upregulation of genes. We only saw upregulation of a handful of genes associated with actin function upon ABP knockdown (**Fig. S9D**), suggesting that actin disruption does not activate a general compensatory upregulation of actin-associated genes. Finally, only mild changes in lipid-related genes were seen across tissues (**Fig. S9E**).

Since the intestine had the largest number of DEGs of any tissue upon ABP knockdown, we next performed GO analysis to further investigate specific pathways that are altered specifically in the intestine upon ABP knockdown. We found that *arx-2* knockdown led to a larger immune response signature, while *unc-60* knockdown had a balance of both immune response and metabolism related changes in gene expression **(Fig. 5D-E)**. On the other hand, *lev-11* resulted in changes completely unrelated to either metabolism or immune response **(Fig. 5F)**. Here, we intentionally opted to avoid overanalyzing changes in neuronal gene expression, as *C. elegans* neurons are generally refractory to RNAi^78^. However, neurons are not entirely refractive to RNAi knockdown and are also amenable to intercommunication effects from other tissues, and thus it is not surprising that many changes were still detected in neurons.

### Targeting of the actin cytoskeleton using small molecule inhibitors impacts aging

Genetic approaches are an excellent method for mechanistic studies, but are more limiting for translational studies, especially in humans. As an alternative, small molecule inhibitors, particularly those isolated from natural sources, offer advantages in delivery and robust downstream effects^79^. Marine natural products have emerged as a rich source of next-generation chemical probes for studying the cytoskeleton^80^, neuroscience^81^, and mitochondrial dynamics^82,83^. To complement our genetic tools, we next employed small-molecule modulators of the actin cytoskeleton. LatA, which sequesters G-actin and promotes filament depolymerization^84^, and Jasp, which stabilizes F-actin and enhances polymerization^85^ to directly destabilize or stabilize the cytoskeleton, respectively. LatA and JASP were purified from dichloromethane repository extracts of *C. mycofijiensis* and *J. splendens* respectively and chemically validated by ^1^H NMR in comparison to literature values^86,87^ (**Fig. S10A-B**).

As expected, we found that destabilization of actin using LatA resulted in a dose-dependent decrease in lifespan (**Fig. 6A**), mirroring our findings using RNAi knockdown. In contrast, Jasp increased lifespans at low concentrations, but decreased lifespans at higher concentrations (**Fig. 6B**), suggesting that increasing stability of actin can be beneficial to a certain degree, after which it can have toxic effects. To validate that treatment with these compounds alter F-actin stability in expected ways, we visualized F-actin using LifeAct::mRuby. As expected, we found that LatA profoundly reduced visible F-actin structures in both the hypodermis and muscle (**Fig. 6C-D, S11A-D**). Interestingly, Jasp increased formation of filamentous actin structures that do not normally exist in the hypodermis, similar to previously reported structures when actin stability was increased via overexpression of the transcriptional regulator, *bet-1*^2^ (**Fig. 6E-F**).

**Fig 6:**
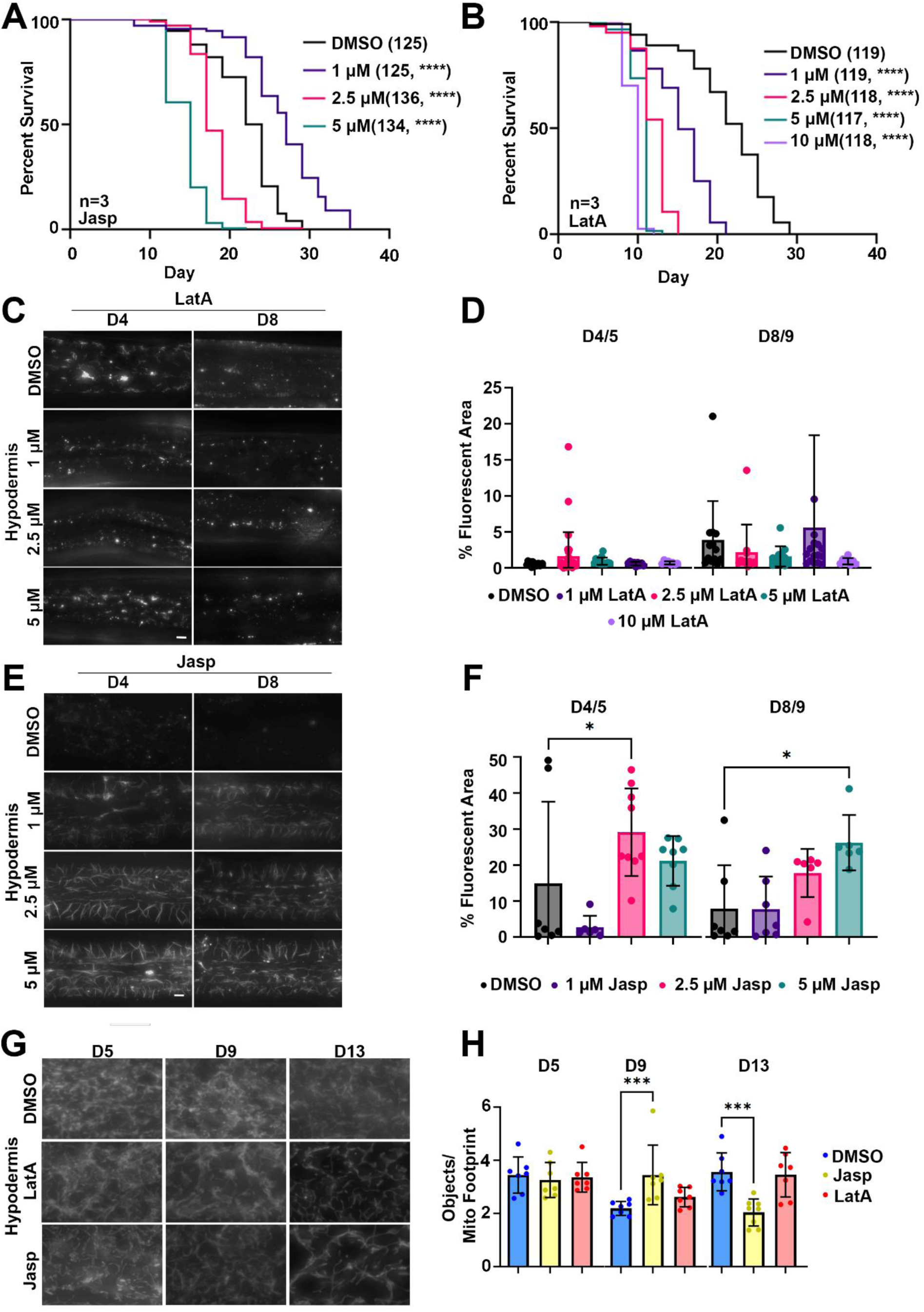
Small molecule actin manipulation alters lifespan and mitochondrial networks. **(A)** Lifespans of wild-type N2 animals grown on empty vector (EV) and varying concentrations of Jasplakinolide (1µM (purple), 2.5 µM (pink), 5 µM (green), 0 µM (control, black)) from day 1 of adulthood. **(B)** Lifespans of wild-type N2 animals grown on empty vector (EV) and varying concentrations of Latrunculin A (1µM (purple), 2.5 µM (pink), 5 µM (green), 10 µM (light purple), 0 µM (control, black)) from day 1 of adulthood. See **Table S1** for lifespan statistics. **(C)** Representative fluorescent images of adult animals expressing LifeAct::mRuby in the hypodermis grown on empty vector (EV) and varying concentrations of Latrunculin A (1µM, 2.5 µM, 5 µM) or DMSO control. **(D)** Quantification of hypodermal actin structures in animals expressing LifeAct::mRuby in the hypodermis grown on empty vector (EV) and varying concentrations of Latrunculin A (1µM, 2.5 µM, 5 µM) or DMSO control. N=3 n>5. **(E)** Representative fluorescent images of adult animals expressing LifeAct::mRuby in the hypodermis grown on empty vector (EV) and varying concentrations of Jasplakinolide (1µM, 2.5 µM, 5 µM) or DMSO control. All animals were imaged on day 4, and 8 of adulthood and images were captured on a Leica Thunder Imager. **(F)** Quantification of hypodermal actin structures in animals expressing LifeAct::mRuby in the hypodermis grown on empty vector (EV) and varying concentrations of Jasplakinolide (1µM, 2.5 µM, 5 µM) or DMSO control. N=3 n>5 **(G)** Animals expressing *col-19p::MLS::GFP* were grown on empty vector (EV), and 1µM Latrunculin A and 1µM Jasplakinolide starting day 1 of adulthood, along with DMSO control, and were imaged during days 5, 9, and 13 of adulthood. **(H)** Quantification was performed using mitoMAPR. Objects/mitochondrial footprint are shown here, and all mitochondrial measurements measured by mitoMAPR are available in **Table S8**. In mitoMAPR-based quantification, the “objects/mitochondrial footprint” refers to the total number of objects detected as mitochondria and the area occupied by all the mitochondria within a defined region of interest. Scale bar for all images is 10 µm.

Similarly, Jasp treatment increased actin filament stability in the muscle at low dose (**S11C-D**). Altogether, our data show that small molecule inhibitors of actin can be robustly utilized for in vivo aging experiments and mirror phenotypes found using genetic approaches.

To further evaluate the effect of Lat and Jasp on actin on downstream cellular processes, we measured mitochondrial morphology in animals treated with 1 µM LatA and Jasp, since they led to opposing results in lifespan. Here, we utilized the hypodermal mitochondria strain, as it exhibits the most profound changes in mitochondrial morphology with age^88^. As expected, treatment with LatA resulted in a qualitative increase in mitochondrial fragmentation in terms of morphology, though this difference did not result in statistical significance when quantified using mitoMAPR (**Fig. 6G-H**). Interestingly, Jasp treatment also resulted in increased fragmentation of mitochondria at late age, despite having displayed increased lifespans. However, considering actin’s critical role in both mitochondrial fusion and fission^89^, it is possible that altering actin dynamics in either direction can perturb mitochondrial morphology.

Finally, we performed bulk RNA sequencing on animals treated with 1 µM LatA and 1 µM Jasp from day 1 of adulthood after 24 hours (Day 2) and 120 (Day 6) hours of drug exposure. RNA-seq libraries showed Phred scores >35 with consistent mapping and clustering across replicates, indicating high-quality data (**Fig. S12**). We found that LatA led to clear changes in the transcriptome after 24 hours of exposure, which was even more pronounced by Day 6 of adulthood (**Fig. S13A-B**). Importantly, almost all DEGs of LatA-treated animals overlapped with those found in *arx-2* and *unc-60* knockdown (**Fig. S13F-K**), providing further evidence that our chemical and genetic approaches have similar effects. However, chemical treatments had significantly fewer DEGs compared to ABP knockdown. Indeed, we found that Jasp had an incredibly mild effect on the transcriptome with no significant DEGs at Day 1 and only 6 significant DEGs at Day 6 of adulthood (**Fig. S13C-D**). Both Jasp and LatA exposure did not significantly alter gene expression of actin-regulatory genes, except for *act-5*, which was significantly increased in expression (**Fig. S13E**). Altogether, these data suggest that the effects of LatA and Jasp mirror some, but not all, phenotypes with genetic perturbations of actin. Moreover, LatA and Jasp likely impact actin and longevity independent of effects on the transcriptome.

### Single-nucleotide polymorphism analysis reveals associations between actin and aging in humans

Based on the incredibly high degree of conservation of actin across eukaryotes and the centrality of its function, we reasoned that actin could also influence aging phenotypes in humans. Here, we focused on genetic variation through naturally occurring single-nucleotide polymorphisms (SNPs) within the ACTB gene locus and measured their potential association with phenotypes corresponding to human aging, with particular emphasis on those related to muscle aging. Here, we analyzed available data of genome-wide association studies (GWAS) in the Health and Retirement Study (HRS); a nationally representative longitudinal study of >36,000 adults over age 50 in the United States^90^. HRS contains biological and genetic samples on subsets of participants and includes physical and psychosocial measures of all study participants in older adulthood, including multiple measures of muscle-related functionality.

SNPs rs852540 and rs73047183 showed suggestive association with a slower rate of decline in gait speed over time in the HRS Non-Hispanic White sample (**Fig. 7A**). Specifically, with each additional G allele of SNP rs852540, there was an average increase in the decline in gait speed of 0.035 meters per second per year compared to other same-aged individuals without the allele (**Fig. 7B**), p-value = 0.0329, surpassing the liberal suggestive cut-off of 0.05). This was assessed among N= 3684 older individuals with a mean of baseline age of 66.33 years (SD=8.26) and a mean of decline in gait speed of -0.002 meters per second (SD=0.0005). We also confirmed that the HRS cohort exhibits an expected age-related decline in gait speed (**Fig. 7C**).

**Fig 7:**
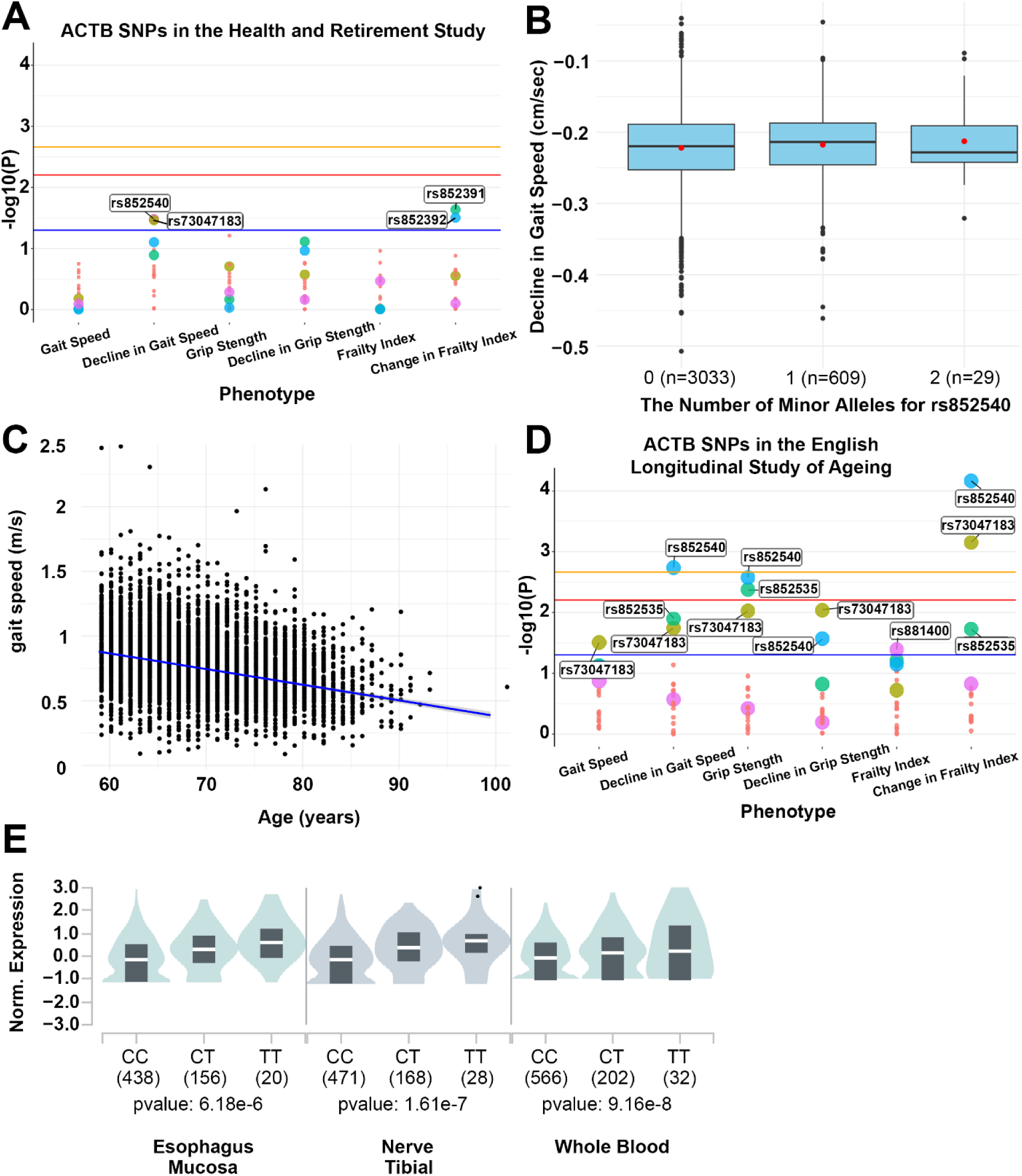
Genome-Wide Association Studies reveal a link between actin and aging outcomes in humans. **(A)** Plot of SNPs associated with age-related phenotypes in human populations from the Health and Retirement Study (HRS). **(B)** The boxplots show the interquartile range for the decline in gait speed in centimeters per second (y-axis) by number of alleles (x-axis) for the SNP rs852540 in ACTB. **(C)** Plot of gait speed by age in HRS. The x-axis shows the baseline age of participants, 50 to 95 years old. The y-axis shows gait speed in meters per second. The sample intercept=0.7968 and slope=-0.00217. **(D)** Plot of SNPs associated with age-related phenotypes in human populations from the English Longitudinal Study of Aging. Blue, red and yellow horizontal lines correspond to p-values of 0.05, 0.00625 and 0.00217, respectively. Each color represents a unique SNP, whereas the large dots represent each SNP that has surpassed the suggestive significance threshold at P<.05 for at least one phenotype. **(E)** Association between rs852535 and ACTB gene expression levels with tissue-specific effects for Esophagus-Mucosa, Nerve-Tibial tissue, and whole blood. Normalized gene expression levels for ACTB (y-axis) by SNP genotype (x-axis) is shown in violin plots. Also on the x-axis is the sample size by genotype in parenthesis. Violin plots show the density plot of the data (green/gray cloud) with the median of the data shown by the white line of the gray box plot within, the lower and upper border of the box plot corresponding to the first and third quartiles, respectively. A linear regression model was used to estimate the mean difference in expression levels, calculated as a Normalized Effect Size (NES) to compare the alternative allele, T, to the minor allele, C.

The two SNPs (rs852540 and rs73047183) showed replication in their effects on decline in gait speed in an independent study cohort, from the English Longitudinal Study of Ageing (ELSA) (**Fig. 7D**), highlighting robust reproducibility across human cohorts. In addition, rs852535 and rs7781531 showed significant associations in ELSA, suggesting that differences also exist across cohorts. Among the four SNPs, rs7781531 showed a faster rate of decline (beta=-0.030) with each additional T allele of rs7781531. This was evaluated among N= 6182 older individuals with a mean baseline age of 66.03 years (SD= 6.79), mean gait speed of 0.77 m/s (SD= 0.29), and the average level of decline in gait speed at -0.015 m/s (SD= 0.01). These effects are examples where variation in the gene contributes to phenotypes that represent different age-related change in functionality; overall we find there are small effects associated with each phenotype, but there are possible pleiotropic effects, and environmental or behavioral factors contributing. It is not known if any one of the identified ACTB SNPs is a causal variant or if they mark a different variant within the ACTB gene that was not represented on the genotyping arrays. Regardless, these results collectively support associations between ACTB and age-related physical function.

We tested replication of the top two SNPs from the GeneWAS across ethnic subsamples in the HRS by calculating a common effect size across the samples. We did this by completing fixed effects and random effects meta-analysis using PLINK software. The Cochrane’s Q statistic (Q), as an indicator of variance across sample effect sizes, and the heterogeneity index (I), which quantifies dispersion across samples, indicate that random effects analysis fit the data better for gait speed decline; thus, we focus on results from random effects to account for differences in effect sizes by sample. The I index indicates 34.84% of the observed variance between samples is due to differences in effect sizes of SNP rs852540, whereas there is 0.0% heterogeneity in the results across studies for rs73047183. The meta-analysis results thus show that only rs73047183 remains associated with decline in gait speed across all race/ethnicity groups (fixed effect p-value=0.0305 and random effect p-value=0.0305). The trait analyzed in the HRS came from a population-based study and was not assessed to allow us to identify physiological degeneration in specific muscles, rather to index and track overall age-related decline in functionality over time. It is widely accepted that genetic variation underlying these aging-related traits are highly polygenic. Thus, it is not expected that a single variant within a gene would be identified to drive these phenotypic results in humans. It is likely that small effects of multiple SNPs across multiple genes, including within the same gene, and with non-additive effects (e.g., gene-by-environment effects), contribute to the resulting phenotypes. With this use of genewide-association scanning approach, it is only possible to identify variants associated with overall effects. Without identifying a causal SNP, we can only aggregate available data to suggest what contributes to a biological pathway. For example, through exploitation of the publicly available Genotype-Tissue Expression (GTEx) database, we found that one of the tag SNPs in ACTB, rs852535, was significantly associated with differential ACTB expression levels through the association with an expression quantitative trait locus (eQTL) in whole blood (p-value=9.16e-8), Esophagus or Mucosa (p-value=6.18e-6), and Nerve or Tibial (p-value=1.61e-7) (**Fig. 7E**).

## Discussion

The actin cytoskeleton is a fundamental player in cellular organization and plasticity, providing the dynamic force and scaffolding for essential processes including vesicular trafficking, mitochondrial dynamics, and protein quality control. While actin dysfunction has been described in disease contexts and associated with several aging contexts, evidence of its direct impact on aging pathology is more limited. Here, we demonstrate that actin disorganization and dysfunction is not simply a consequence of aging but can be causative of the onset of aging phenotypes and exacerbate multiple hallmarks of aging, which directly correlate with a reduction in lifespan. We found that genetic disruption of actin or ABPs resulted in an expected acceleration of age-associated decline in actin structures, which has direct physiological outcomes including a reduction of lifespan, impaired motility, and reproductive decline (**Table 1**). Perhaps most critical in our findings is that each ABP exerted distinct effects in each tissue: *unc-60* knockdown compromised intestinal actin organization as early as day 1 of adulthood, *lev-11* knockdown impaired muscle filaments and contractility, and *arx-2* disruption broadly perturbed filament networks across tissues. These phenotypic outcomes directly mirrored their effects on lifespan. Thus, our studies highlight the key importance of each ABP uniquely in specific tissues. These results add to the current understanding that actin is not a homogeneous structure but a collection of specialized assemblies that fulfill unique functions in different tissues. Therefore, our findings help to uncover potentially novel methods to target actin for therapeutics by focusing on unique cell-type specific requirements of ABPs, without perturbing actin structure or function universally across the animal, which can have profoundly detrimental off-target effects.

**Table 1.** Summary of the effect of actin disruption on healthspan and lifespan.

| Phenotype\RNAi |  | *act-1* | *arx-2* | *unc-60* | *lev-11* |
| --- | --- | --- | --- | --- | --- |
| Lifespan |  | ↓↓↓ | ↓↓↓ | ↓↓↓ | ↓↓↓ |
| Motility |  | ↓↓↓ | ↓↓↓ | ↓↓↓ | ↓↓↓ |
| Brood |  | ↓↓↓ | ↓↓↓ | ↓↓↓ | ↓↓↓ |
| Mitochondrial Fragmentation | Muscle | ↑ | ↑↑↑↑ | ↑ | ↓ |
|  | Hypodermis | ↑ | ↑↑ | ↑ | ↓ |
|  | Intestine | ↑ | ↑ | ↑↑ | ↓ |
| Respiration (OCR) |  | ↓ | ↓↓↓ | ↓↓↓ | ↓↓↓ |
| Lipid droplets | DHS-3::GFP signal | ↓ | ↓ | ↓↓↓↓ | ↓↓↓↓ |
|  | Droplet number | ↑ | ↑ | ↑↑↑ | ↑ |
|  | Droplet size | ↓↓ | ↓↓↓ | ↓↓ | ↓ |
| Protein aggregation | Neuron | ↑ | NC | ↑↑↑ | ↑↑↑ |
| Autophagy (LGG-1 puncta) | Muscle | ↓↓↓↓ | ↓↓ | ↓↓↓↓ | NC |
|  | Hypodermis | NC | NC | NC | NC |
|  | Intestine | ↑↑↑↑ | ↑↑↑ | ↑↑↑↑ | ↓↓↓ |
| Gut barrier integrity |  | ↓ | ↓ | ↓ | ↓ |

NC: No Change; ↓/↑: trend; ↓↓/↑↑: p<0.05; ↓↓↓/↑↑↑: p<0.01; ↓↓↓↓/↑↑↑↑: p<0.001

Beyond phenotypic profiling, we performed transcriptomic profiling, which revealed that actin perturbations broadly induce an aged signature, with acceleration of transcriptomic age most pronounced in *arx-2* and *unc-60* knockdowns. Although DEGs varied across conditions, immune and defense response genes were consistently upregulated in all conditions, in line with actin’s established role in maintaining epithelial junctions to protect from immune invasion. This is an important consideration for *C. elegans* biology, as the bacterial food source is ultimately the cause of death upon breakdown of the gut barrier^91–93^. In the wild, *C. elegans* have a mixed bacterial diet and are constantly exposed to bacteria of various pathogenicity. Therefore, regulation of proper immune responses and host-pathogen interaction are critical for animal survival. Further studies are necessary to determine whether actin-related changes to the gut barrier and expression of immune-related genes directly translate to effects on pathogen resistance.

Mitochondrial dysfunction is one of the most well-established hallmarks of aging across species, and perhaps one of the most heavily studied roles of actin in relation to aging. For example, actin is required for asymmetric inheritance of mitochondria during budding in *S. cerevisiae* to create a pristine daughter cell with a full replicative lifespan^50^. Moreover, increased actin polymerization in the aging fly brain results in excessive F-actin accumulation, which impairs autophagy and increased mitochondrial dysfunction to shorten lifespan^10^. Although direct effects of actin-mitochondria interactions on longevity have not yet been revealed in mammalian systems, considering the well-established role for actin in mitochondrial dynamics^89^ and direct correlation of dysregulation of mitochondrial fusion/fission dynamics on numerous metrics of aging pathology^94–101^, it is not a stretch to consider direct impacts of age-associated actin dysregulation on cellular and organismal health directly through its impact on mitochondrial dynamics. Indeed, we find that directed actin dysfunction results in major changes to mitochondrial structure and function. Importantly, altering actin in any direction (either increasing or decreasing stabilization using small molecules) resulted in increased mitochondrial fragmentation, which is a critical consideration for targeting actin for mitochondrial dynamics.

Since many of the mitochondrial fusion and fission factors are shared, including the actin cytoskeleton^89,102,103^, it is challenging to predict the outcome of how alterations in actin can affect mitochondria. For example, our data show that genetic knockdown of actin, *arx-2*, or *unc-60* resulted in fragmentation of mitochondria, but knockdown of *lev-11* resulted in increased fusion. Since tropomyosins bind to actin, which can affect both stability^104,105^ and myosin motor binding to actin to impact contractility^106–108^, it is possible that disruption of tropomyosin impinges on both fusion and fission machinery dependent on both processes. Disruption of both fusion and fission generally has been shown to a retain relatively tubular network^109^. Thus, it is plausible that tropomyosin knockdown, by simultaneously impairing fusion and fission to varying degrees, might shift the mitochondrial network toward a hyperfused state.

Beyond just general mitochondrial function, stress responsive transcription factors are also shared between the actin cytoskeleton and mitochondria. For example, the heat-shock transcription factor, HSF-1, has been shown to be important for maintenance of actin stability with age^9^ and in regulation of the UPR^MT 110,111^. In our data, we found that knockdown of actin and ABPs also directly impact gene expression of mitochondrial stress responsive genes.

Finally, increased integrin signaling results in increased stability of actin, remodeling of mitochondria, and activation of the mitochondrial stress response, which can have direct effects on longevity in *C. elegans*^112^ and potentially impact cancer aggression in mammalian models^113^. Altogether, our data adds to the increasing amount of circumstantial evidence linking an actin-mitochondria axis directly to aging pathology. Lipid homeostasis was also impaired, with actin and ABP knockdown animals displaying an increased number of significantly smaller lipid droplets. Although lipid dysregulation is often concomitant with mitochondrial dysfunction, there is usually an increase in lipid content due to impairment of beta oxidation of lipids through the mitochondria^114,115^. Thus, in animals with actin dysfunction, it is possible that mitochondrial dysfunction results in a compensatory increase in lipid metabolic pathways to supply fatty acids for energy. Indeed, we find that these animals exhibit a significant upregulation of lipid metabolic genes. Moreover, lipids can be cleared and processed via autophagic turnover^116,117^, and indeed, *arx-2* and *unc-60* animals displayed an increase in autophagosome puncta in the intestine. This is also consistent with a reduction in polyQ in the intestine, which can also be due to increased clearance into autophagosomes^118^. However, considering actin’s critical role in autophagosome fusion to lysosomes and its subsequent clearance, it is also possible that the increase in autophagosomes could result from failure in completion of the late stages of autophagy, and future work is critical to determine whether the increase in autophagosomes upon *arx-2* and *unc-60* knockdown is due to a compensatory increase in autophagy, a failure in clearance of autophagosomes, or possibly a mix of both.

Importantly, changes in autophagy were not universal across all actin dysfunction conditions or across tissues. Interestingly, *lev-11* knockdown displayed a reduction in autophagosome puncta in the intestine, while displaying increased autophagosomes in the muscle. Considering *lev-11* encodes a tropomyosin primarily expressed in the muscle, it is interesting that there are impacts of *lev-11* knockdown across several phenotypes in non-muscle cells. This could potentially be due to non-autonomous signals from the muscle that impact non-muscle cells. For example, muscle-specific overexpression of TFEB, one of the primary transcription factors that regulate autophagy, can result in improved brain function through muscle-derived chemokines, often called “myokines”^119^. Finally, muscle-specific activation of the transcription factor FOXO can result in organism-wide activation of proteostasis pathways and improve longevity^120^. Thus, it is entirely feasible that even upon damage, the muscle can release non-autonomous signals to enact the function of other cell-types. However, why or how muscle-defects can induce defects in other tissue remain to be explored. One tantalizing hypothesis is that damaged muscle cells may attempt to restore homeostasis by jettisoning damage to other tissues, a phenomenon commonly found in neurons^62,121^. For example, muscle cells can secrete aggregated prion proteins to adjacent muscle tissue, and even to the hypodermis, a process dependent on autophagic machinery^122^. Thus, upon *lev-11* knockdown, it is possible that muscle cells attempt to ameliorate cellular dysfunction by transferring damaged components to other tissues. An entirely different hypothesis is that *lev-11* is not completely a muscle-specific gene, and that either undetectable or stress-induced expression of *lev-11* has some important functions in non-muscle cells.

Pharmacological perturbations reinforced that actin dynamics must be finely tuned for healthy aging. Reducing actin stability with LatA resulted in an expected shortening of lifespan, while stabilization with Jasp extended lifespan at low doses but became deleterious at higher concentrations. This is likely due to the beneficial effect of mildly stabilizing actin filaments; however, hyper-stabilizing of actin can become toxic due to suppression of dynamic and fluid functions of actin. This was reminiscent of increased actin stabilization via *bet-1* overexpression. In specific tissues, like the muscle and intestine, where increased actin stability was beneficial for their function, *bet-1* overexpression extended lifespan. However, in the hypodermis where dynamic actin is likely required for endocytosis, exocytosis, and signaling events, *bet-1* overexpression resulted in ectopic expression of actin fibers, which reduced lifespan^2^.

Interestingly, both LatA and Jasp increased mitochondrial fragmentation, highlighting the intricate and complex regulation of actin dynamics on mitochondria. This data also underscores that changes to mitochondrial morphology alone do not predict functional outcomes on lifespan when actin remodeling is perturbed. Consistent with this, *lev-11* knockdown resulted in a hyperfused mitochondrial network, despite also reducing lifespan.

The lack of large-scale transcriptomic changes following drug treatment suggests that in these conditions, actin primarily influences aging through structural and post-transcriptional mechanisms rather than broad transcriptional reprogramming. Important to note, LatA and Jasp were administered to animals starting at day 1 of adulthood due to developmental lethality when applied during larval stages. In opposition, RNAi conditions were applied from hatch. Thus, cellular and physiological changes to actin are likely influenced both by non-transcriptional and transcriptional changes downstream of actin dysfunction. If applied during development, actin dysfunction potentially results in major shifts in chromatin remodeling – which actin has increasingly become known to regulate^123–125^ – thus driving major transcriptional changes throughout the lifespan. In opposition, if applied after development, actin dysfunction only has mild impacts on transcriptome and thus exerts its effects more directly on actin structure and organization. This is similar to other stress paradigms, such as mitochondrial hormesis, which has been shown to be required to be applied during larval stages^126^, which results in dramatic chromatin remodeling to elicit effects throughout the lifespan^127^.

The evolutionary conservation of actin’s role in cellular processes is well established, especially considering the immense sequence and structural similarity of actin across all eukaryotes^1^. Considering the shared roles of actin in numerous cellular and physiological processes across organisms, we predicted that actin’s potential role in aging is similarly conserved. Here we found that common polymorphisms in ACTB were associated with differences in gait speed decline across two independent aging cohorts, suggesting that natural variation in actin regulation contributes to functional trajectories in older adults. Although effect sizes were modest, replication across cohorts strengthens the conclusion that changes in actin can influence neuromuscular aging in humans.

### Limitations of the study

In keeping within the scope of a resource, we deliberately present our findings in a data-forward manner and refrain from overinterpreting our data. Rather than forcing a single tone or conclusion, we contextualize our data within the broader literature without making unwarranted mechanistic claims, thus providing a resource that offers a platform for future investigators to mine, integrate, and build upon.

While our study presents evidence that actin structure and function play a key role in the aging process, several limitations should be noted for future studies. First, GWAS associations that we found are correlational in nature, and the identified SNPs were found in noncoding regions. Therefore, testing causality or molecular mechanisms of these SNPs driving aging pathology is challenging, as these noncoding regions are not shared in model organisms, like *C. elegans*. Another limitation is that the human cohorts analyzed are predominantly of European ancestry, raising questions about generalizability of our findings and the potential differences in other populations.

In our *C. elegans* studies, RNAi was applied from the L1 stage of development, raising the possibility that developmental perturbations contributed to adult phenotypes. In addition, the RNAi method itself has several limitations: first, RNAi knockdown efficiency varies across tissues. This is particularly true where some tissues – like the pharynx or neurons – are known to be refractory to RNAi^70,128,129^. This is further exacerbated by the complex genetics required for cell-type specific RNAi, which requires mutation and tissue-specific rescue of RNAi machinery. Since these RNAi machineries have important canonical roles in gene expression and pathogen defense^130–132^ – there are likely physiological consequences of knockout and subsequent overexpression in cell types that naturally do not have such high expression of these genes. To complicate things further, many of the genes encoding ABPs have multiple isoforms or gene variants that could serve redundant roles, often with specific cell-types expressing each gene in variable levels. However, the dramatic rewiring of the transcriptome at day 1 suggests that effects during development are likely to have lasting effects throughout adulthood. Moreover, our confirmation that knockdown of *arx-2, lev-11,* and *unc-60* robustly perturbed actin organization in expected ways provide confidence that redundant genes were not compensating for their loss. Finally, our adult-specific chemical approaches replicating many phenotypic outcomes of RNAi suggest that not all effects of actin dysfunction on organismal health and longevity are through developmental effects and that our RNAi conditions are highlighting outcomes of direct actin perturbations. However, more precisely controlled manipulations in future work will help separate developmental versus adult-specific effects, as well as better understand the timing of when actin interventions can be applied during the life course to impact physiology and longevity.

One limitation of using *C. elegans* as a model system as a whole is that they are post-mitotic during adulthood, meaning our findings may not fully capture how actin dysfunction can impact highly proliferative tissues, such as in the liver or skin of mammals. This is an important consideration since one of actin’s critical roles is in cell division, and numerous studies implicate changes to actin dynamics in both hyperproliferative cancer cells and non-proliferative senescent cells^133–137^. Thus, further work in animal models with proliferative cells throughout the adult life course is essential to further translate our findings.

Finally, our work focuses primarily on end-point measurements of actin: that is, how filament structure, organization, and function impact cellular and organismal physiology. However, actin is a complex multi-protein structure made up of the collective function of hundreds to thousands of actin subunits interacting with dozens to hundreds of actin binding proteins. Therefore, future work will need to focus on how changes to actin or ABP translation, folding, post-translational modification, and interaction of these thousands of proteins define the end-point measurements of actin filament assembly, organization, and function. Together, these caveats highlight both the challenges and the opportunities in defining actin as a key player in organismal health and longevity. Nevertheless, the convergence of genetic, pharmacological, and human GWAS data argues that actin integrity is indispensable for maintaining cellular health across aging tissues, and that preserving actin homeostasis may represent a powerful strategy for manipulating organismal healthspan.

## Author Contributions

M.A. designed and performed all *C. elegans* experiments and prepared the manuscript and figures. H.N., J.K., S.M.G., M.C.W., and G.G. assisted with transcriptomics analysis. T.W., A.A., T.N., D.S., M.V., N.D., R.A.B., A.J.C., D.M., T.C.T., M.O., M.P., and S.P.C. assisted with *C. elegans* lifespans and imaging experiments. C.M. and C.K. assisted with autophagy experiments. J.A.G., S.F.O., J.G.C., N.S., M.S.N., J.C.P., H.L-T, and T.A.J. purified LatA and Jasp. H.M. performed seahorse assays. M.A.T. assisted with biochemistry. E.M.C. and T.E.A. performed all GWAS analysis. G.G. and R.H.S. assisted with manuscript preparation and figure construction, designed, executed, and analyzed all experiments and data.

## Data Availability

All data required to evaluate the conclusions in this manuscript are available within the manuscript and Supplementary Materials. All strains synthesized in this manuscript are derivatives of N2 or other strains from CGC and are either made available on CGC or available upon request. All raw RNA-seq datasets are available through Annotare 2.0 Array Express Accession E-MTAB-16193 and E-MTAB-16195. All code used to analyze datasets included in this work are available on the Sanabria lab GitHub repository: https://github.com/SanabriaLab/Averbukh-et-al.-2026

All the GWAS phenotype and genotype data are available to approved researchers. HRS: https://www.ncbi.nlm.nih.gov/projects/gap/cgi-bin/study.cgi?study_id=phs000428.v2.p2 ELSA: https://www.ebi.ac.uk/ega/studies/EGAS00001001036

## Acknowledgements

M.A. and A.J.C. is supported by T32AG052374 from the NIA; T.W. is supported by the COMPASS CIRM grant; M.V., M.O., and T.C.T. are supported by 1R25AG076400 from the NIA; J.K. is supported by the USC Provost Fellowship; E.M.C. is supported by P30AG068345; S.P.C. is supported by NIH R01AG058610 and Hevolution Foundation award 748 HF AGE-004; T.A.J. is supported by The Fletcher Jones Endowment fund of DUC; M.C.W. is supported by the Howard Hughes Medical Institute; T.E.A. is supported by P30AG068345; C.K. is supported by NIA grant R01AG083373; G.G. is supported by supported by T32AG052374 and R01AG079806-02S1 from the NIA; and R.H.S. is supported by R00AG065200 and R01AG079806 from the NIA, the Glenn Foundation for Medical Research and AFAR Grant for Junior Faculty Award, and 2022-A-010-SUP from the Larry L. Hillblom Foundation. Some strains were provided by the CGC, which is funded by the NIH Office of Research Infrastructure Programs (P40 OD010440). GWAS analysis was performed at the Gerontology Bioinformatics Core at the University of Southern California and was supported by the University of Southern California and Buck Institute Nathan Shock Center P30AG068345. Single-nuclei RNA-sequencing was performed with the assistance of the Visiting Scientist Program at the HHMI Janelia Research Campus. Real-time oxygen consumption rates (OCR) of *C. elegans* were measured using the XF96 Analyzer (Seahorse Bioscience) at the USC Leonard Davis School of Gerontology Seahorse Core. We thank Dr. Hasan Celik (UC Berkeley) and Pines Magnetic Resonance Center’s Core NMR Facility (PMRC Core) for spectroscopic assistance with the instruments used in this work that were supported in part by the PMRC Core and NIH S10OD024998.

The HRS (Health and Retirement Study) is sponsored by the National Institute on Aging (grant number NIA U01AG009740) and is conducted by the University of Michigan.

The ELSA (English Longitudinal Study of Ageing) is sponsored by the National Institute of Aging [grants 2RO1AG7644-01A1 and 2RO1AG017644] and a consortium of UK government departments coordinated by the Office for National Statistics. Funding for the English Longitudinal Study of Ageing genotyping data was provided by the Economic and Social Research Council (ESRC) and by the National Institute of Aging grants.

The content is solely the responsibility of the authors and does not necessarily represent the official views of the National Institutes of Health. The funders had no role in study design, data collection and analysis, decision to publish, or preparation of the manuscript.

## Materials and Method

### C. elegans strains and maintenance

All strains used in this study are derived from the N2 wild-type strain from the Caenorhabditis Genetics Center (CGC). All *C. elegans* strains are maintained at 15 °C on Nematode Growth Media (NGM) plates containing 1 mM CaCl2, 12.93 μM (5 µg/mL) cholesterol, 25 mM KPO_4_ (pH 6.0), 1 mM MgSO_4_, 2.5 mM (0.25% w/v) peptone, 51.3 mM NaCl, and 2% w/v agar. Plates are seeded with a saturated OP50 *E. coli* B strain grown in standard Luria-Bertani (LB) medium at room temperature for 24-48 h. Animals are synchronized using a standard bleaching protocol^27^ where worms are collected in M9 buffer (22 mM KH₂PO₄, 42.3 mM Na₂HPO₄, 85.6 mM NaCl, 1 mM MgSO₄), then transferred into a 1.8% sodium hypochlorite and 0.375 M NaOH suspension. Animals are vigorously shaken in sodium hypochlorite solution until only eggs remain, which are then pelleted using centrifugation at 1,100 x g and washed with M9 solution four times. Eggs are then floated in M9 overnight (up to 16 h) at 20 °C on a rotator to synchronize at the L1 stage and plated onto RNAi plates containing 1 mM CaCl2, 12.93 μM (5 µg/mL) cholesterol, 25 mM KPO_4_ (pH 6.0), 1 mM MgSO_4_, 2.5 mM (0.25% w/v) peptone, 51.3 mM NaCl, 1 µM IPTG, 100 µg/ml carbenicillin, and 2% w/v agar (Note: we strictly use BD Difco granulated agar (VWR, 90000-782) as different agar brands can have different substrate stiffnesses that affect *C. elegans* physiology^138^). RNAi plates are seeded with HT115 *E. coli* K strain, which carry either the pL4440 empty vector (EV) or a pL4440 construct expressing double-stranded RNA against a gene of interest (all conditions described in figure legends and all RNAi sequences used in this study are available below). Animals are grown at 20 °C for all experiments unless otherwise noted. Day 1 adult animals are defined as gravid adult animals, meaning animals with a full egg sac, which occurs approximately 3 days (72 h) from the L1 stage at 20 °C.

*act-1* RNAi

TGTGTGACGACGAGGTTGCCGCTCTTGTTGTAGACAATGGATCCGGAATGTGCAAGGCCGGATTCGCCGGAGACGACGCTCCACGCGCCGTGTTCCCATCCATTGTCGGAAGACCACGTCATCAAGGAGTCATGGTCGGTATGGGACAGAAGGACTCGTACGTCGGAGACGAGGCCCAATCCAAGAGAGGTAAATAATTAATACATTCGATGATTAAATTTATGCGTACTATTTCAGGTATCCTTACCCTCAAGTACCCAATTGAGCACGGTATCGTCACCAACTGGGATGATATGGAGAAGATCTGGCATCACACCTTCTACAATGAGCTTCGTGTTGCCCCAGAAGAGCACCCAGTCCTCCTCACTGAAGCCCCACTCAATCCAAAGGCTAACCGTGAAAAGATGACCCAAATCATGTTCGAGACCTTCAACACCCCAGCCATGTATGTCGCCATCCAAGCTGTCCTCTCCCTCTACGCTTCCGGACGTACCACCGGAGTCGTCCTCGACTCTGGAGATGGTGTCACCCACACCGTCCCAATCTACGAAGGATATGCCCTCCCACACGCCATCCTCCGTCTTGACTTGGCTGGACGTGATCTTACTGATTACCTCATGAAGATCCTTACCGAGCGTGGTTACTCTTTCACCACCACCGCTGAGCGTGAAATCGTCCGTGACATCAAGGAGAAGCTCTGCTACGTCGCCCTCGACTTCGAGCAAGAAATGGCCACCGCCGCTTCTTCCTCTTCCCTCGAGAAGTCCTACGAACTTCCTGACGGACAAGTCATCACCGTCGGAAACGAACGTTTCCGTTGCCCAGAGGCTATGTTCCAGCCATCCTTCTTGGGTATGGAGTCCGCCGGAATCCACGAGACTTCTTACAACTCCATCATGAAGTGCGACATTGATATCCGTAAGGACTTGTACGCCAACACTGTTCTTTCCGGAGGAACCACCATGTACCCAGGAATTGCTGATCGTATGC

*arx-2* RNAi

TTTCAGGCGAAATGGATTCGCAAGGGCGAAAGGTGATTGTCGTTGACAACGGAACAGGTGTAAGTTGCAAAATTTTATCCCATTTTATTTTCAAAAATTCTGTATTTGCAGTTCGTCAAATGCGGGTATGCAGGAACCAATTTCCCAGCTCATATATTCCCTTCAATGGTCGGTCGGCCAATCGTGAGATCGACACAAAGAGTTGGAAATATTGAAATTAAGGTAAAACTTTACACAGAAAGGTCTTATAACTGTTTTCTCACAGGATTTGATGGTTGGCGAGGAGTGCTCTCAGCTTCGTCAAATGCTTGATATCAATTATCCCATGGACAACGGAATTGTTAGAAACTGGGATGATATGGCGCATGTATGGGATCATACTTTTGGACCGGAAAAGTTGGATATCGACCCGAAAGAGTGTAAACTGCTGCTGACAGAACCACCACTCAATCCGAACAGCAATCGTGAGAAAATGTTTCAAGTTATGTTTGAGCAGTATGGATTCAATTCTATCTATGTGGCGGTTCAGGTAATAGTTAATCCTCATGATAAAATCATAATGATTATATTTTCAGGCTGTGCTCACTCTTTATGCTCAAGGTCTTTTGACAGGAGTTGTAGTTGACTCCGGAGATGGTGTTACCCACATCTGTCCTGTTTACGAAGGATTTGCTCTTCATCATTTGACAAGACGATTGGATATTGCAGGAAGAGATATTACCAAGTATCTTATCAAGGTAAGTTTTTAAATGTTACATATTAATTCAATTATTTTCCTGATTGCTATAGTGACAAAATATG GTTTATTAGAAACTAACATATATTTGTTTTCAGCTTCTTCTGCAACGTGGATACAACTTCAATCACTCTGCTGACTTTGAAAC

*lev-11* RNAi

TGTCGAAGGTAAACAAGGAGGGAGCTCAGCAGACATCGCTTCTTGATGTCCTCAAGAAGAAGATGCGTCAGGCCCGTGAGGAGGCTGAGGCCGCCAAGGACGAGGCCGACGAGGTCAAAAGACAGCTCGAGGAGGAGCGCAAGAAGCGTGAAGATGTATGTGATTATAGTAGCGACGACTAGGTTTCTTGTGATAAGTCTCAATAATTCCATCTAGTGTAACAAGCCCCGCGCGGGCACTAAGTCAATATAAGTTGTATAAAGTGGCTGCTCCTGCTGCTTATTTGTCTTAGAATAGGAAAAAGTGATGCGAGAGAAAGTGAATGAGGCTATGGGATTTTGTAGGCTGAAGCCGAGGTTGCCGCCTTGAACCGTCGTATTGTGCTTGTGGAGGAGGACTTGGAAAGAACCGAAGATCGTCTGAAGACCGCCACATCAAAGCTTGAACAAGCAACCAAAGCTGCCGATGAGGCTGATCGGTTCGTTTTTTTCTATCCCATTTGTGTCAATTACAGGCTGAGGCTGAGGTCGCTTCTTTGAACCGTCGCATGACCCTCCTCGAGGAAGAGCTCGAGAGAGCCGAGGAGCGTCTGAAGATCGCCACCGAGAAGCTCGAAGAGGCTACCCACAATGTCGACGAGTCTGAGCGGTAGGCGTTGAGATGAGGATTCCTGTGGATCTCAAGCCAAACTACCAAACTACTGCCAGTGCTCAGAAAATTATATATATATTGTGAACGTCATTGTCCCAATTGTGTACTGTAAATTTTTTAGCGCGCGCAAGTCGATGGAAACCCGCTCCCAACAGGACGAGGAACGTGCCAACTTCCTCGAGACTCAAGTCGACGAGGCTAAGGTTATCGCCGAGGATGCTGATCGCAAATACGAAGAGGTGCCGATTGAATGCGCGTGCTTAATGGCTTAATGGCTTAATGTTAACTGTCATGTGTCCTGAATGTTAAACCCCCCTAATAGTTTGGAGTTTTTTTTTTACAAAATCTAATCTCTATTATAGATCCAGAAGAGCGTTGAGCAATCAGATTGACATGGATGATGACAGATGTTCGGATCTTGAAAGGAAGTTAAGAGAGTGTCAGTCGATTTTGCATGAAACTGAAAACAAAGCAGAAGAGGTAGAGCTTTATTTACTTTCAACTCCTAGAAATTCCATGTTCTAATGCTTTATGTTGTCCGAGTTGTCCGAACACTGGCACAAGAATACAATGGCACAGAGATCCTATCAGGCGATCCCTCACCCCGAAACTTTTGTTGAATAATACTGTTCCAGCGTGCGCAAGGTTATGGAGAACCGCTCCCTTCAGGATGAGGAGCGCGCCAACACCGTTGAGGCCCAACTTAAGGAGGCTCAACTTCTCGCCGAGGAGGCTGACCGCAAATACGACGAGGTTACCATCGAGTGGATTATATATATATATATTATATAATCATATAACATGTTCTCATACTAATTGTCTTGCATCTTTAACACTATAACATTAGACGTATGTGACAAAAATAATAATGAAAATTGTTCCAGGTCGCCCGTAAGCTCGCCATGGTTGAAGCTGATCTTGAGAGAGCTGAGGAGCGTGCCGAGGCCGGAGAGAAGTGAGTTAAGAGCACCTATCACAATGCCCCCTTTTCATTTACTTTTACGCTTCTTAACCCAACTAACAAAAACGCATAATTCTAATAAAACCGATAAAACATAAAACCACCATCACAGCAAGATCGTGGAGCTTGAAGAGGAGTTGCGCGTCGTTGGTAATAACTTGAAATCCCTTGAACTTTCCGAGGAAAAGGCACTTGAGAAGGAGGACATCTTTGCCGAGCAGATTCGTCAGCTCGATTTCAGACTGAAGGAGGTATCAACGTGTCTCAGAAAGTCTCTCTTCACCTCAAAACCTTCCGGAATACCTTTTGAGATGATATGCTTATTAACAAACTCATTCGCTTTAACAATCAAGCTTCAAACTTTGTATTCCTCTAATCTTAACTTGGGTGACATTCTTAGCCTAACCTTACAGACGATCTAACCTGGCAGAGGCCCACATGCGCGGGCTCTCCGTGAATCTTCGTGAGGCGCAGGACCTGTTACATCAGCTGCAGCAGGAGGAGAACGATTCCTGCGAATACTTGAATTGCGCCGTAGAATCGCGAAAGGAGGTTTCTCAAACACCATTTGTTCATAAAACTAATGCTTCTGCACGACACTAAACTCTTACTCATTCCCATCTTCACATCTCTCTCTCTCCCCCTTTCAAACCGTCGTTGTCCCATCTTTCAGCAAGATCGTCGAACTCGAGGAGGAGCTCCGTGTCGTCGGAAACAACTTGAAGTCACTTGAGGTGTCCGAGGAGAAGGCTCTCCAACGTGAGGACTCGTACGAGGAGCAGATCCGCACCGTCTCATCCAGACTGAAGGAGGTCCGATTCTTTTCTGTGTAGTAGTCTTTCTGTGTCTTCACTATCAAAGTGATGTCCGTGTTGTCTTTAATTGTGATAATAGTTCCTGAAAAAAAAACGAAATTTCTTTCAGGCTGAGACCCGTGCCGAATTCGCTGAGCGTTCCGTCCAGAAGCTCCAGAAGGAGGTCGACAGACTCGAAGGTAGGTCTAAGGGGCTTTATATGTGATATCTAAATGTCATTGTCAATTTGTGTAAGTCTATATGTGTATTGTCCACGTGTTCAGATGAACTCCTCCTCGAGAAGGAGCGTGTCCGAAACTTGACTGAAGAAATCGAACAGACCGTCCAGGAGATCCAAGGATCCTA

*unc-60* RNAi

ATGGTGAGTTTGAGATTTTAATTCGCTTCAATATTTTTAAATTGAAAGAAATTTGGAATGTTA TGTTTGGCGTGTTTTTTTTTCATTGAATTTCCAAAATTTCACACTAAAATGGAATGGTTTTCT CTTCTTTTTGCCTAACCTAACCTAACCTATGTGTGCCTGTTTTCTAGAGTTCCGGTGTCATG GTCGACCCAGATGTGCAGACATCTTTCCAAAAGCTCTCCGAGGGACGCAAGGAGTACCGC TACATCATTTTCAAGATCGACGTGAGTTTTTAAAAAATAAAATCTGAATCAGATCAATTTAAA AAAATTTCGTGCCACTTTTTGTTTTTTTTGTTGAAAAATTTGAAAAATCCTCAAATTAATTGTT TTAGATGAACATTATTGATTTCCCTATTAAATTGCAACATTTTCCAGGAGAACAAGGTGATC GTGGAAGCCGCGGTGACTCAGGATCAGCTCGGCATCACTGGAGACGACTATGATGACTCT TCCAAGGCCGCTTTCGACAAATTCGTCGAGGACGTGAAGTCTCGAACCGATAATCTGACC GATTGCCGCTACGCCGTTTTCGACTTCAAGTTCACGTGCAGTCGTGTTGGAGCCGGCACG AGCAAGATGGACAAGATCATCTTCCTCCAGATGTAAGCGCTTGATCCTATTGGTGAATTATT GTACCATCTAATTTTTTTTCCAGCTGCCCAGATGGTGCTTCTATCAAGAAAAAGATGGTGTA CGCTTCGTCCGCCGCCGCCATCAAGACTTCTCTCGGAACCGGCAAAATCCTTCAGTTCCA GGTGAGAAATCTCGATAATTTTTACAATTAGAAAAAAAAAATCAAATTATTAAAATTTCAGGT GTCTGACGAGAGCGAGATGAGCCACAAGGAACTCCTCAACAAGTTGGGCGAGAAATACGG AGATCACTAG

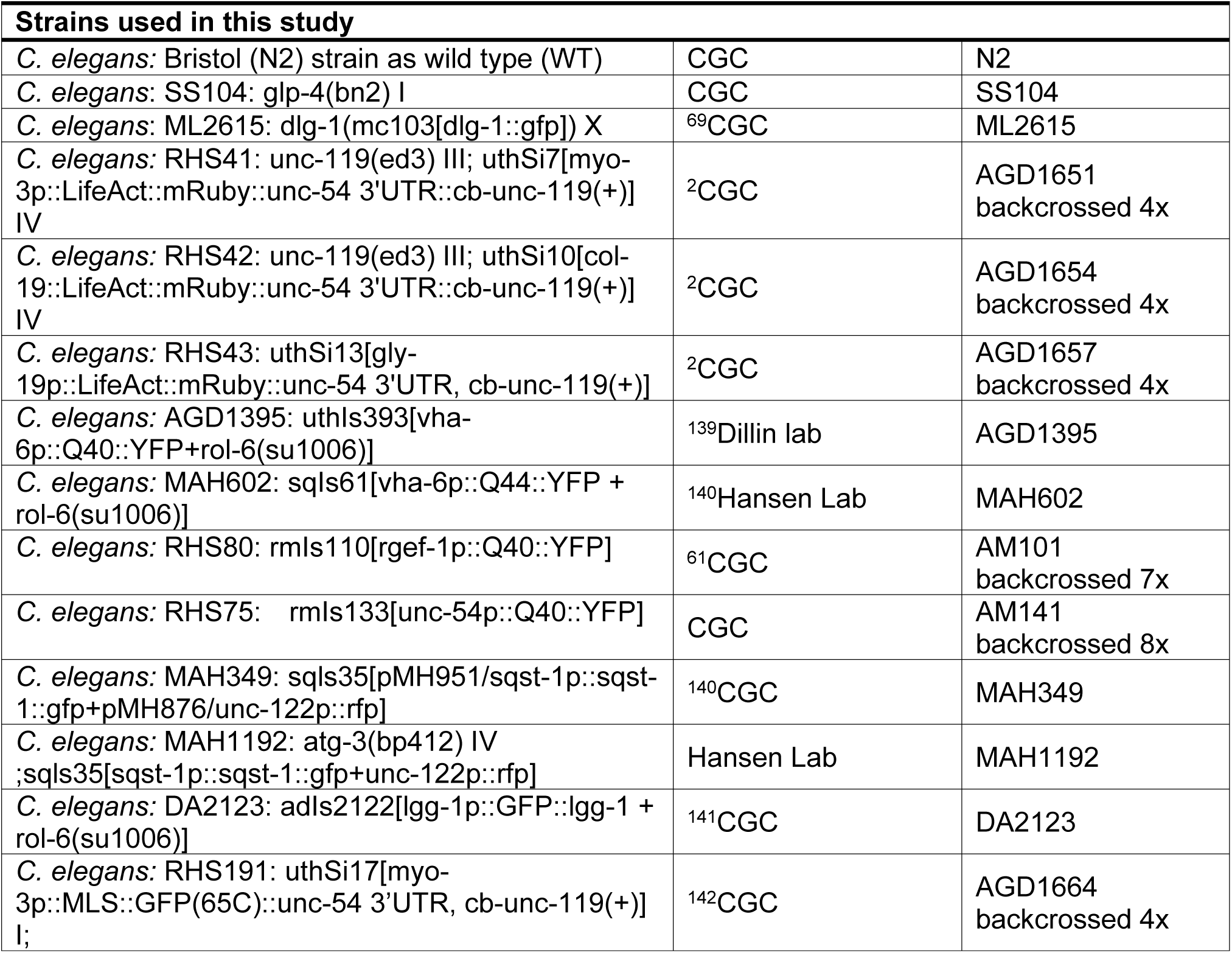
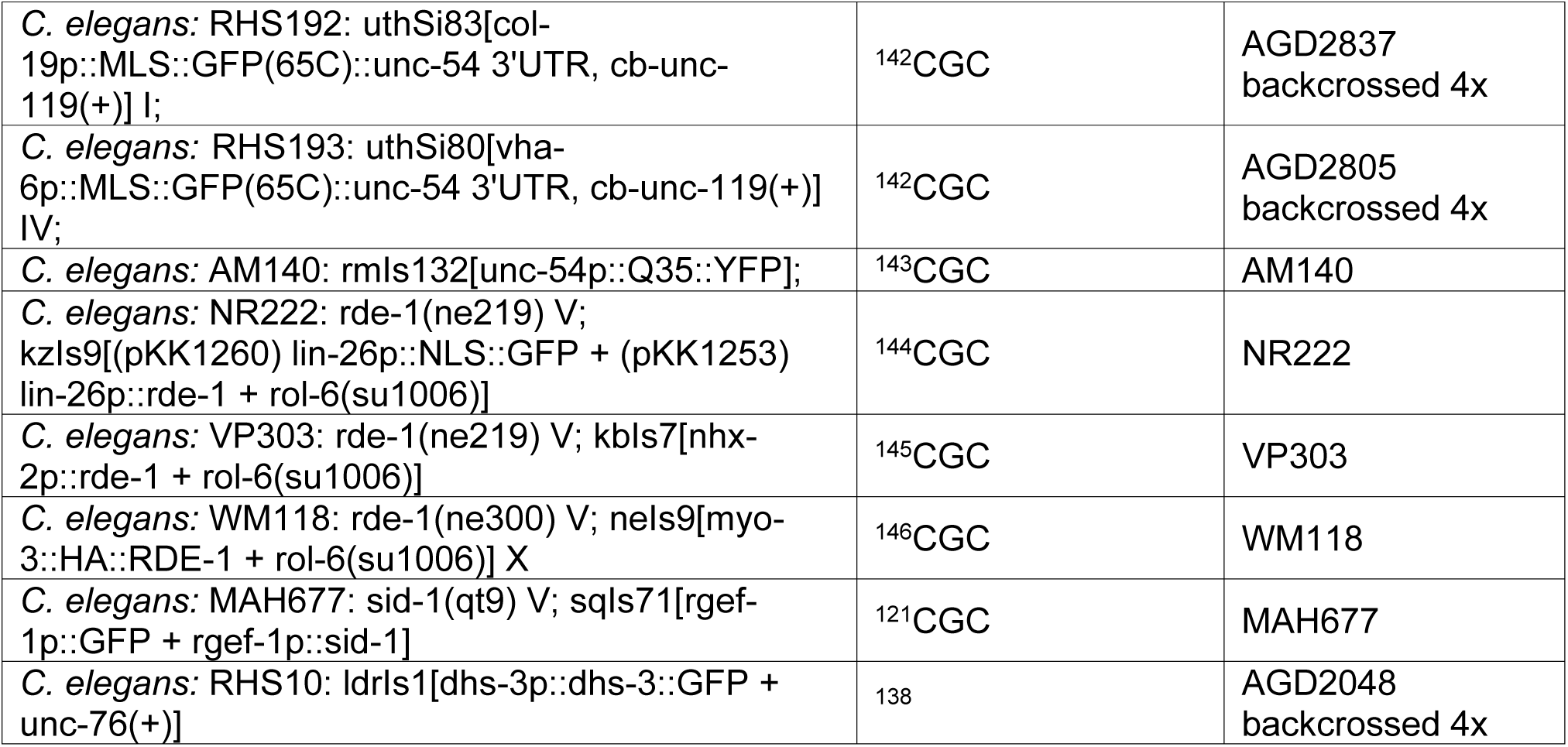

All high-resolution microscopy was performed on live animals at the specified ages in the figures and figure legends. Animals were synchronized using a standard bleaching protocol and all animals were aged on RNAi plates supplemented with FUDR starting from day 1 of adulthood until the desired age as previously described^147^. To make FUDR plates, 100 µL of 10 mg/mL FUDR is spotted onto an RNAi plate seeded with RNAi bacteria of choice. Bacteria are allowed to grow for a minimum of 24 h on RNAi plates prior to seeding with FUDR. For imaging, animals are picked from plates and mounted directly onto a microscope slide into M9 + 0.1 M sodium azide to anesthetize animals. For standard wide-field imaging, imaging was performed on a Leica Thunder Imager equipped with a 63x/1.4 Plan ApoChromat objective, standard dsRed filter (11525309) and GFP filter (11525314), a Leica DFC9000 GT camera, a Leica LED5 light source, and run on LAS X software. For confocal microscopy, imaging was performed on a Stellaris 5 confocal microscope equipped with a white light laser source and 405 nm laser source using spectral filters, HyD detectors, 63x/1.4 Plan ApoChromat objective, and run on LAS X software. For all microscopy, representative images of three independent biological replicates are shown.

For full-body imaging of *C. elegans*, animals are lined up on NGM plates without bacteria. ∼10-12 worms are placed into M9 + 0.1 M sodium azide to anesthetize worms, arranged in a line with the pharynx facing up, and imaged with a Leica M205FCA automated fluorescent stereomicroscope driven by LAS X software. The microscope is equipped with a standard GFP and RFP filter and a Leica K5 camera.

For quantification of muscle actin, the number of muscle cells with defective actin filaments were counted manually in each animal. Muscle cells were scored as having defective actin if any visible filaments were shown to display any observable differences in phenotype (e.g., fraying of filaments, uneven filaments, broken filaments, etc.). The % of muscle cells displaying disrupted actin vs. total number of muscle cells were calculated per image and data is presented as each biological replicate plotted as a single data point. Statistical analysis was performed on GraphPad Prism using parametric student t-testing. Intestinal actin structures were evaluated by blinded manual scoring on a scale of 1–3, where 1 indicated normal wild-type–like morphology, 2 indicated partially disturbed structures, and 3 indicated severely disrupted structures

For quantification of mitochondrial morphology, we used mitoMAPR^52^, which quantifies numerous metrics of mitochondrial morphology. Here, we used mitochondrial object length and measurement of object number/mitochondrial footprint as representative measurements.

Mitochondrial object length is generally higher for fused mitochondria and lower for fragmented mitochondria. Mitochondrial object number/mitochondrial footprint measures the total number of distinct mitochondrial objects per volume of mitochondria and higher numbers indicate more fragmented mitochondria while lower numbers indicate more fused mitochondria.

Hypodermal actin structures were quantified in Fiji by calculating the percentage of fluorescent area relative to the total image area (**Source Code 1A**). Generally, this macro first clears the “Results” table, enables Batch Mode for faster, window-less processing (by stopping images from flashing on the screen), and sets measurement fields once (area, mean, min, area_fraction, 3 decimals). It then loops over all open images (nImages), selecting each in turn and calling processImage. Inside processImage, it notes where new rows will start (prevNumResults) and grabs the image title for labeling, switches to channel 1 (note this depends on how many channels are imaged, but this should be the optimal fluorescent channel), converts the image to 8-bit, and sets an auto-threshold (“Triangle white”). Measure records intensity stats over the entire image (so mean/min come from the grayscale image), while area_fraction is computed from the current threshold (fraction of pixels above threshold). After measuring, it writes the filename into the newly added rows. Once all images are processed, the macro turns Batch Mode off, refreshes the Results table, prompts for a save location, and saves the table with area, mean, min, area_fraction calculation for each image.

DHS-3::GFP signal in stereoscope images was quantified in Fiji by drawing an ROI around each animal and measuring the integrated fluorescence intensity (**Source Code 1B**). This macro quantifies lipid droplet signal by looping over all currently open images and measuring user-drawn ROIs. It first closes any existing Results table, initializes one by setting measurements (area, mean, integrated density, 3 decimals) and making a dummy measurement, then clears it so it’s ready. For each image (nImages), it selects the image, switches to channel 2, converts to 8-bit, and lightly enhances contrast (0.35% saturation). It resets the ROI Manager, pauses to let the user draw ROIs around each worm and add them to the ROI Manager, then iterates through those ROIs. For each ROI it runs Measure (using the current selected channel), tags the new row in Results with a label like Worm_#_<IMAGETITLE>, and updates the table. After all images/ROIs are processed, it asks where to save and writes the Results table as “IntegratedIntensity_Summary_(rep #).csv” in chosen folder by the user.

Gut assay images were quantified in Fiji by drawing an ROI around each animal and measuring the integrated fluorescence intensity above a threshold (**Source Code 1C**). This macro measures fluorescent signal in the gut on channel 2 for every open image, using only pixels above a set threshold inside user-drawn ROIs. It first closes any existing Results table, initializes measurement settings (area, mean, integrated density, 3 decimals) with “Limit to threshold” enabled, makes a dummy measurement to create the table, then clears it. For each image (nImages), it selects the image, switches to channel 2, converts to 8-bit (so the threshold is on a 0–255 scale), and prompts the user to draw ROIs around each worm and add them to the ROI Manager. For each ROI, it reselects channel 2, applies a fixed threshold of 25–255 (so only brighter pixels count), keeps “Limit to threshold” active, and runs Measure, producing Area (of the ROI above threshold), Mean (mean intensity of those included pixels), and Integrated Density (Area × Mean) for that worm. It tags each row with a label Worm_<#>_<IMAGETITLE>, updates the Results table, and after all images/ROIs are processed, asks where to save and writes the CSV as “day#_rep#_IntegratedIntensity_Summary_gut_assay.csv”.

Neuronal polyQ foci were quantified in Fiji by drawing a region of interest (ROI) around each individual animal and measuring both the number of aggregates and the size of each aggregate (**Source Code 1D**). Similarly, DHS-3::GFP puncta in confocal images were quantified in Fiji by measuring both the number and size of lipid droplets per image and using the full image as the ROI (**Source Code 1D**). In short, this macro loops over every open image, switches to channel 3 (can be whichever channel contains the fluorescent images for analysis), and asks the user to draw ROIs (around each worm for stereoscope images and the full frame for confocal images) in the ROI Manager. For each ROI, it duplicates just that region into a new image and runs “Clear Outside” so only the area of interest remains. It then preprocesses the cutout: converts to 8-bit, does rolling-ball background subtraction (radius 30 px) to remove smooth background, applies MaxEntropy auto-threshold to segment bright signal, sets black background, converts to a binary mask, and runs a quick erode (dilate to clean up noise and fill small gaps). Next, it sets measurements and calls Analyze Particles… with “summarize”, which populates the Summary table with the usual totals (e.g., Count, Total Area, Average Size, %Area) for that ROI region. It then renames the “Slice” entry in the Summary table to a label like Worm_#_<IMAGETITLE> so you can trace results back to each worm/image, close the cutout, and continue to the next ROI/image. When finished, it prompts for a folder and saves the Summary table as “Counts_SummaryTable.csv”.

### C. elegans bulk RNA-seq

RNA isolation was performed at day 1 of adulthood in *glp-4(bn2)* animals grown at 22 °C to ensure that no progeny or gametes contribute to transcriptional changes. This was imperative as we show in this manuscript that perturbing actin function can have dramatic effects on reproduction, and thus removing progeny is critical for whole-animal assays. ∼1000 animals were used to harvest RNA. Specifically, worms were washed off plates with M9 and immediately placed in Trizol solution. RNA isolation was performed by freeze/thaw cycles in liquid nitrogen/37 °C bead bath cycles 3x. After the final thaw, chloroform was added at 1:5 chloroform:trizol ratio and the aqueous separation was collected via centrifugation in a heavy gel phase-lock tube (VWR, 10847-802). The aqueous phase was mixed 1:1 with isopropanol, then applied to a QuantaBio Extracta Plus RNA kit (76492-576) column and RNA purification was performed as per manufacturer’s directions.

For bulk RNA-seq, library construction and sequencing were performed at Novogene on an Illumina NovaSeq6000 using a poly(A) selection, first-strand synthesis, and paired-end workflow. Three biological replicates were analyzed for each condition. Low quality reads and adaptor sequences were trimmed using Trim Galore v0.6.7-1^148^. Reads were aligned to the WBcel235 genome using STAR-2.7.3a^149^, and a raw count matrix was generated using Subread v2.0.3^150^. Differential gene expression was calculated with DESeq2^151^ and sva v3.54.0 was used to correct for batch effects and non-biological sources of variation^152^. GO enrichment analysis was run through WormEnrichr^153,154^.

The Upset plots were plotted using the R package UpSetR version 1.4.0^155^. Fisher’s exact test was performed using the ggbarstat() function of the R package ggstatsplot version 0.13.1^156,157^. For predicting biological age, we first converted raw counts into count per million (CPM). We then computed predicted biological age using elastic net coefficients and the code provided in the BiT age publication^35^.

### Worm preparation for nuclear isolation

Conditional germline-less mutant *glp-4(bn2)* worms grown at permissive 15 °C until gravid adult stage. Eggs were isolated through the established bleach-based protocols. Eggs were allowed to synchronize to L1 overnight in M9 at 20 °C and then plated onto 60 mm RNAi plates containing 10X concentrated EV or RNAi bacteria at a concentration of 250 worms per plate.

The worms were grown at the restrictive temperature of 22 °C until day 1 of adulthood before being collected for nuclear isolation. An estimated 3,000 worms were collected per condition.

### Nuclear isolation

All buffers were made fresh prior to worm collection and placed on ice. All tips, tubes, and pestles were rinsed with homogenization buffer or PBS with BSA to reduced adhesion of nuclei to the surfaces. Collected worms were washed 3x with M9 in a 15 mL tube and then transferred to a 1.5 mL tube. The tubes were spun down in a swinging bucket centrifuge at 4 °C and the supernatant was removed before the samples were placed on ice. 100 μL of homogenization buffer was added to the worm and then they were ground with a motorized pestle for 45 seconds. 800 μL of homogenization buffer was added to the samples while used to wash the pestle. The samples were transferred to sterile 1 ml Dounce homogenizers on ice (Wheaton, 357538). 20 strokes were applied to the samples using the loose pestle and 50 μL of homogenization buffer were used to rinse the pestle into the sample. 20 strokes of the tight pestle were then applied and 50 μL of homogenization buffer were once again used to rinse the pestle into the sample. The lysates were then filtered into a 35 μM cell strainer tube, followed by a Flowmi cell strainer (Bel-Art, H13680-0040) to transfer the lysate to a 1.5 mL tube. The filtered lysates were spun at 800 xg for 10 minutes at 4 °C using a swinging bucket rotor. The supernatant was removed and the pellet was resuspended in 500 μL of PBS with 0.5% BSA and RNAase inhibitor by pipetting 60 times and avoiding the formation of bubbles. Once the pellet was fully resuspended, the samples were filtered once more through a Flowmi cell strainer while being transferred to a flow cytometry tube. 20 μL of the filtered samples was transferred into a tube containing 180 μL of PBS with 0.5% BSA and RNAase inhibitor to be used as an unstained control for Fluorescence Activated Cell Sorting (FACS). The remaining sample was stained on ice using Hoechst (Invitrogen, 33342) at a working concentration of 1:1,000. Approximately 200,000 to 300,000 Hoechst positive events were sorted based on forward scatter area (FSC-A) into a 1.5 mL tube. Sorted nuclei were spun at 1,000 xg for 10 minutes at 4 °C using a swing bucket rotor. The supernatant was removed and the nuclei were resuspended in 45 μL of PBS with 0.5% BSA and RNAase inhibitor by pipetting. 2 μL of the resuspended nuclei were diluted and counted using a hemocytometer based on Hoechst fluorescence.

### snRNA-seq

18 μL at a concentration of 1,000 nuclei/μL were provided to the Janelia Research Campus Quantitative Genomics core and run through the 10X Genomics Chromium Single Cell 5’ (v2 Chemistry Dual Index) pipeline. scRNA-sequencing was performed by Novogene using their NovaSeq XPlus 25B Premade-10X 5 prime Single Cell Transcriptome Library pipeline.

### snRNA-seq pre-processing

Raw reads were processed using Cell Ranger (9.0.0, 10X Genomics). Sequencing reads were aligned to a Cell Ranger protein coding reference generated using *C. elegans* genome WBcel235.dna.toplevel.fa and the WBcel235.113.gtf files. Outputted raw and filtered Feature Barcode Matrices were loaded into R (4.4.3) using Seurat (4.4.0) and processed based on previously published workflow^158^. Matrices were decontaminated using SoupX (1.6.2) and converted to Seurat objects (SeuratObject_4.1.4) with thresholds set to a minimum of 3 cells and minimum of 100 features. Outliers were filtered by subsetting for 100-8000 features, less than 5% mitochondrial genes, and less than 20,000 counts. Normalized data was scaled with SCTransform (0.4.1) and doublets were removed with DoubletFinder (2.0.3) and SCDS (1.22.0) R packages using the proportion estimated by 10X Genomics user guide (CG000204 Rev D).

### snRNA-seq clustering and cell-type annotation

Singlets were scaled using SCTransform with variable features set to 5000 and nFeature_RNA and percent.mito as variables to regress. Samples were integrated using 2000 variables features selected using vst method. Integration anchors were identified using rpca reduction and k.anchor set to 20. Dimensional reductions using Uniform Manifold Approximation and Projection (UMAP) methods were performed using 50 dimensions from Principal Component Analysis. Clusters were identified using default Louvain algorithm, 100 random starts, and a 0.8 resolution. Cell type annotations were assigned using the SingleR package (2.8.0). and canonical cell types annotated in a previous snRNA-seq dataset as a reference^159^. Clusters were then annotated as the tissue making up more than 50% of the cluster or based on established worm cell type proportions using the two largest cell types within the cluster.

### Bulk RNA-seq deconvolution and differential expression analysis

Raw bulk RNAseq counts were deconvolved using the InstaPrism package (0.1.6) and snRNAseq data acquired using same strain and conditions as the bulk RNAseq^160^. Cell type specific gene expression counts were deconvolved using the Instaprism get_Z_array() function. Deconvolved cell type specific gene counts were rounded and differential expression analysis was performed using the DEseq2 package (1.46.0). SVA (3.54.0) and lima (3.62.2) packages were employed to correct for non-biological sources of variation and batch effects^161,162^.

### C. elegans thrashing assay

Thrashing assays were performed at specific ages specified in the figure legends by flooding a plate containing adult animals with 100 µL M9 solution. 30 s videos were captured on a Leica M205FCA stereomicroscope using a Leica K5 camera. Trashing was measured by eye over a 10 second period and a single thrash was defined as bending of a minimum of 50% of the animal’s body in the opposite direction. Data is representative of three independent biological replicates as a dot plot generated using Prism 7 software. All statistics were performed using non-parametric Mann-Whitney testing.

### C. elegans brood size assay

Brood assays were performed by moving 10 L4 animals onto individual plates seeded with EV culture. Every 12 hours, animals were moved onto a new plate and the original plate containing eggs were stored at 15 °C for 2-3 days until live progeny were counted and scored. All animals across all progeny plates were summed together per individual animal to determine brood size. Data is representative of three independent biological replicates as a dot plot generated using Prism 7 software. All statistics were performed using non-parametric Mann-Whitney testing.

### C. elegans lifespan assay

Lifespan assays were performed on standard RNAi plates at 20 °C. Day 1 adult animals were placed onto RNAi plates containing FUDR and scored every day for viability. Animals were scored dead if they showed no signs of moving after prodding both the head and the tail with a platinum wire. Animals were scored as censored if they exhibited age-unrelated death including bagging (vivipary), intestinal leaking out of the vulva, crawling up the sides of the wall, burrowing into agar, etc. All statistical tests were performed using Prism7 software using LogRank testing and all statistics are made available in **Table S1**.

For LatA and Jasp lifespans, animals are grown on standard RNAi plates from L1 and at the day 1 adult stage, animals are moved onto RNAi plates containing compounds. LatA and Jasp are added directly to NGM RNAi plates to concentrations indicated in figure legends. DMSO was used as a vehicle control. Lifespans were scored every 2 days similar to standard lifespans.

### C. elegans seahorse assay

Day 1 adult animals were washed off plates with M9 and transferred onto an NGM plate with no bacteria. 10-15 worms were then pipetted off from this plate into individual wells of a Seahorse XF96 microplate, ensuring no progeny were included. Basal oxygen consumption rate was measured using a Seahorse XFe96 Analyzer equipped with an XFe96 sensor cartridge using 2 minutes of mixing, 30 second waiting period, and 2 minutes measurement. A total of 6 measurements were made. Animals were then exposed to 50 mM sodium azide to measure non-mitochondrial respiration rates as previously described^163^. A total of 6 measurements were made after addition of sodium azide using 2 minutes of mixing, 30 second waiting period, and 2 minutes measurement. OCR were plotted using Prism7 software and statistics were performed using one-way ANOVA using GraphPad Prism.

### C. elegans autophagy measurements

Autophagy was monitored by quantifying GFP::LGG-1/Atg8 puncta in body-wall muscle, proximal intestinal cells, and hypodermal seam cells of strains DA2123 and RD202 on day one of adulthood. Animals were grown at 20 °C from hatching on RNAi plates containing EV, *act-1* (1:9 culture with matched OD600), *arx-2*, *unc-60*, or *lev-11*. For imaging and puncta quantification, animals were mounted on 3% agarose pads in M9 medium supplemented with 0.1% sodium azide. GFP::LGG-1/Atg8 puncta were visualized using a Zeiss Imager Z1 equipped with an apotome.2, a Hamamatsu Orca Flash 4LT camera, and Zen 2.3 software.

Puncta were counted in body-wall muscle cells, the two to three most proximal intestinal cells, or one to three visible hypodermal seam cell at 1000× magnification. At least 7-10 animals were imaged per condition, with results pooled from three independent experiments and analyzed using Student’s *t*-test (GraphPad Prism).

### Chemical isolation and purity of latrunculin A (Lat A)

Biological material collection and identification of specimens of the marine sponge *C. mycofijiensis* were collected via scuba diving in Vanuatu, as previously reported^164,165^. Taxonomic identification was conducted by comparing characteristic biological features with reference samples from the UC Santa Cruz sponge repository. Voucher specimens and underwater photographs can be provided upon request. Extracts of *C. mycofijiensis* were processed according to previously reported methods^166–168^. The fat dichloromethane extracts (FD) were used in the repeated scaleup HPLC isolation of pure Lat A. HPLC purification was performed on a semi-preparative column (Phenomenex Inc. Luna© 5μm C18(2) 100 Å 10 × 250 mm) in conjunction with a 4.0 × 3.0 mm C18 (octadecyl) guard column and cartridge (holder part number: KJ0-4282, cartridge part number: AJ0-4287, Phenomenex Inc., Torrance, CA, USA). A reversed-phased linear gradient was employed (30:70 CH_3_CN/H_2_O to 80:20 over 50 min, 100% CH_3_CN from 51 to 61 min, then 30:70 for re-equilibration from 62 to 73 min). Compound detection was measured by means of a single wavelength (λ max = 230 nM) using a Spectroflow 783 programmable absorbance detector (detector part number: 9000-7831). LatA would begin to elute at approximately 52 minutes. The purified compound was dried under an N_2_ gas stream, stored in clear vials, and sealed in a dark desiccator under vacuum. The purity of the compound was confirmed to be >95% by ^1^H NMR analysis using a Bruker 500MHz Avance NEO spectrometer equipped with a Prodigy broadband observe cryoprobe optimized for X-detection

### Chemical isolation and purity of jasplakinolide (Jasp)

Specimens of the marine sponge *J. splendens* (coll. no. 00101) were collected in Fiji, and taxonomic identification were conducted as previously reported^169^. Voucher specimens and underwater photographs can be provided upon request. Samples of *J. splendens* were processed from repository extracts according to previously reported methods^87,169^. The fat dichloromethane extracts (FD) were used in repeated scaleup HPLC isolation of pure jasplakinolide. HPLC was performed on a preparative column (Phenomenex Inc. Luna© 10 µm PREP C18(2) 100 Å 250 x 21.2 mm) in conjunction with a 10.0 × 10.0 mm C18 guard column and cartridge (holder part number: AJ0-9281, cartridge part number: AJ0-7221, Phenomenex Inc., Torrance, CA, USA). A reversed-phased linear gradient was utilized (50:50 CH_3_CN/H_2_O to 100:00 over 30 min, 100% CH_3_CN from 31 to 38 min, then 50:50 for re-equilibrium from 39 to 48 min). Compound detection was measured by means of a single wavelength (λ max = 230 nM) using a Spectroflow 783 programmable absorbance detector (detector part number: 9000-7831). Jasplakinolide elution began at approximately 36 min. The purified compound was dried under an N2 gas stream and stored in clear vials. The purity of the compound was confirmed to be >95% by ^1^H NMR analysis using a Bruker 500MHz Avance NEO spectrometer equipped with a Prodigy broadband observe cryoprobe optimized for X-detection.

### Study Population

Independent discovery and replication samples were used in this study. The discovery sample was drawn from the U.S. Health and Retirement Study (HRS^90,170^), which is a nationally representative, longitudinal sample of adults aged 50 years and older in the US, who were interviewed every two years, beginning in 1992. HRS includes households across the country with over 36,000 participants contributing. The GeneWAS subsample for decline in gait speed was restricted to 3,684 respondents (59.58% female) who (1) aged ≥ 50 years; (2) had at least two completed gait speed measurements; (3) speed ≤2.5 m/s; (4) identified as Non-Hispanic White to eliminate the issues of population stratification; and (5) had available genetic information. The replication samples were drawn from the English Longitudinal Study of Ageing (ELSA), designed as a sister study to HRS. It is a nationally representative study cohort of individuals living in England aged over 50, for which ethical approval and written consent were granted by the London Multi-Centre Research Ethics Committee. This study was initiated in 2002 with 11,391 participants and has since included follow-up every two years. The sample inclusion criteria were the same as HRS, with 6,182 ELSA participants (54.08% female) included in the current study, from whom gait speed was assessed at least two times between 2002 and 2018.

### GWAS Genotyping Data

Genotype data were accessed from the National Center for Biotechnology Information Genotypes and Phenotypes Database (dbGaP^171^). DNA samples from HRS participants included in the present study were collected in two waves (2006, 2008) and genotyped by the NIH Center for Inherited Disease Research (CIDR; Johns Hopkins University) using the HumanOmni2.5 arrays from Illumina (San Diego, CA). Raw data from both phases were clustered and called together. HRS followed standard quality control recommendations to exclude samples and markers that obtained questionable data, including CIDR technical filters^172^ removing SNPs that were duplicates, had missing call rates ≥ 2%, > 4 discordant calls, > 1 Mendelian error, deviations from Hardy-Weinberg equilibrium (at p-value < 10-4 in European samples), and sex differences in allelic frequency ≥ 0.2). Further detail is provided in HRS documentation^173^. Applying these criteria to the gene region, on chromosome 7, (NC_000001.10): 5,566,040-5,603,533 resulted in available data on 18 SNPs with minor allele frequency equal or greater than 0.01 within the ACTB region that are on the Illumina array to represent 240 human SNPs in the gene. With the goal of evaluating whether representative marker SNPs across the gene are associated with the phenotypes of interest, and the significant statistical evidence between associations of ACTB variants and decline in gait speed, a clumping procedure was performed. The procedure selects SNPs by taking the most significant SNPs within the region and prunes variants with redundant correlated effects caused by linkage disequilibrium (LD). To inspect the SNPs relations, SNPs with a minor allele frequency lower than 0.01 among Non-Hispanic White were first excluded, followed by testing the R-squared by applying the parameters of: physical distance threshold at 250 kb, LD threshold of 0.5, and p-value thresholds for index and clumped SNP at the level of 0.98. We then selected the most significant SNP for each group of correlated SNPs, resulting in 12 SNPs (bold SNPs in **Table S2**).

The genotype data in ELSA were performed by University College London (UCL) Genomics Institute using the HumanOmni2.5–8 array from Illumina (San Diego, CA). DNA was extracted from blood samples collected in 2004 and 2008, Because the two studies were designed to be similar (with 2,368,902 markers in common across the studies), the same quality control procedures used with HRS were applied to the ELSA data, except that UCL excluded SNPs with missing call rates >5% (rather than 2%), yielding available data on 1,372,240 SNPs for 7,412 individuals with European ancestry, of whom 6,182 had phenotypic data also.

### Phenotype Construction

Phenotype construction was completed to calculate common measures of normal age-related muscle decline in functionality over time. Phenotypes were calculated after merging multiple survey years to get repeated assessments on the same individuals, and based on consensus in the literature on population-based surveys of aging^174–176^. Gait speed was assessed as the number of seconds taken to walk a 98.5-inch (250 cm) course. Individuals were asked to complete two timed walks (to one end, stop, and back). Gait speed was conducted with respondents aged 65 years or older in HRS and the average of two-timed walks was calculated then converted into units of meters per second (baseline mean = 0.79, SD=0.25). The phenotype for gait speed decline was assessed as slopes, interpreted as changes in performance on gait speed over time. Slopes were calculated using gait speed measures from 2006 to 2018 and a mixed effects model in SAS 9.4, adjusted for sex, age at the first assessment point, and number of years of follow-up. Identical procedures were carried out with ELSA data.

### GeneWAS

GeneWAS was run as a series of linear regression scans, under an additive model, adjusting for relevant covariates and indicators of population stratification. Population stratification occurs when the correlation between dependent and independent variables differs for subpopulations and may result in spurious genetic associations^177^. To reduce such type 1 error, we conducted GeneWAS adjusting for population substructure as indicated by latent factors from principal components analysis (PCA)^178,179^. Detailed descriptions of the processes employed for running PCA are provided by HRS and outlined by Patterson and colleagues^179^. From PCA, the first two eigenvalues with the highest values accounted for less than 4.5% of the overall genetic variance, with additional components (3-8) increasing this minimally, by a total of ∼1.0%^173^.

Based on these analyses, we opted for a strategy that does not ignore population substructure, but also does not over-correct, and adjusted for the first four PCs in all analyses with HRS data. With the ancestral homogeneity in the ELSA dataset, no PCs were used. All GeneWAS were completed using PLINK 2.0^180^.

### SNP Evaluation

We evaluated SNP associations in the GeneWAS by p-value. With the number of SNPs and primary phenotypes in this study, strict Bonferroni correction would yield an adjusted multiple test-correction p-value threshold of 0.0028 (for 18 tests). However, Bonferroni correction such as these are too conservative because of the correlations among SNPs^181,182^ and the cross-validation approach. To address the correlation among SNPs, we implement a clumping schema and calculate empirical p-value thresholds, through permutation^181–184^. Permutation is a process whereby necessary correlations between SNPs and phenotypes are intentionally shuffled so that p-values for the shuffled (null) data are compared to the non-shuffled data. This permutation is repeated multiple times in order to determine an empirical p-value^182,184,185^, a calculated threshold at which a test result is less likely to achieve significance by chance alone. Thus, when performing 10,000 permutations using PLINK and max(T) option, the p-value thresholds of 0.0319 for gait speed decline were observed for determining gene-wide significance. For SNP comparisons, we used R^186^.

### Gene Expression

The Genotype-Tissue Expression (GTEx) database^187^, the most comprehensive, publicly-available resource for tissue-specific gene expression data, was used to evaluate whether there was evidence for regulatory functions of SNPs within the gene. We entered the top SNPs into GTEx to assess relationships with differential gene expression.

## Supplemental Figures

**Fig S1.**
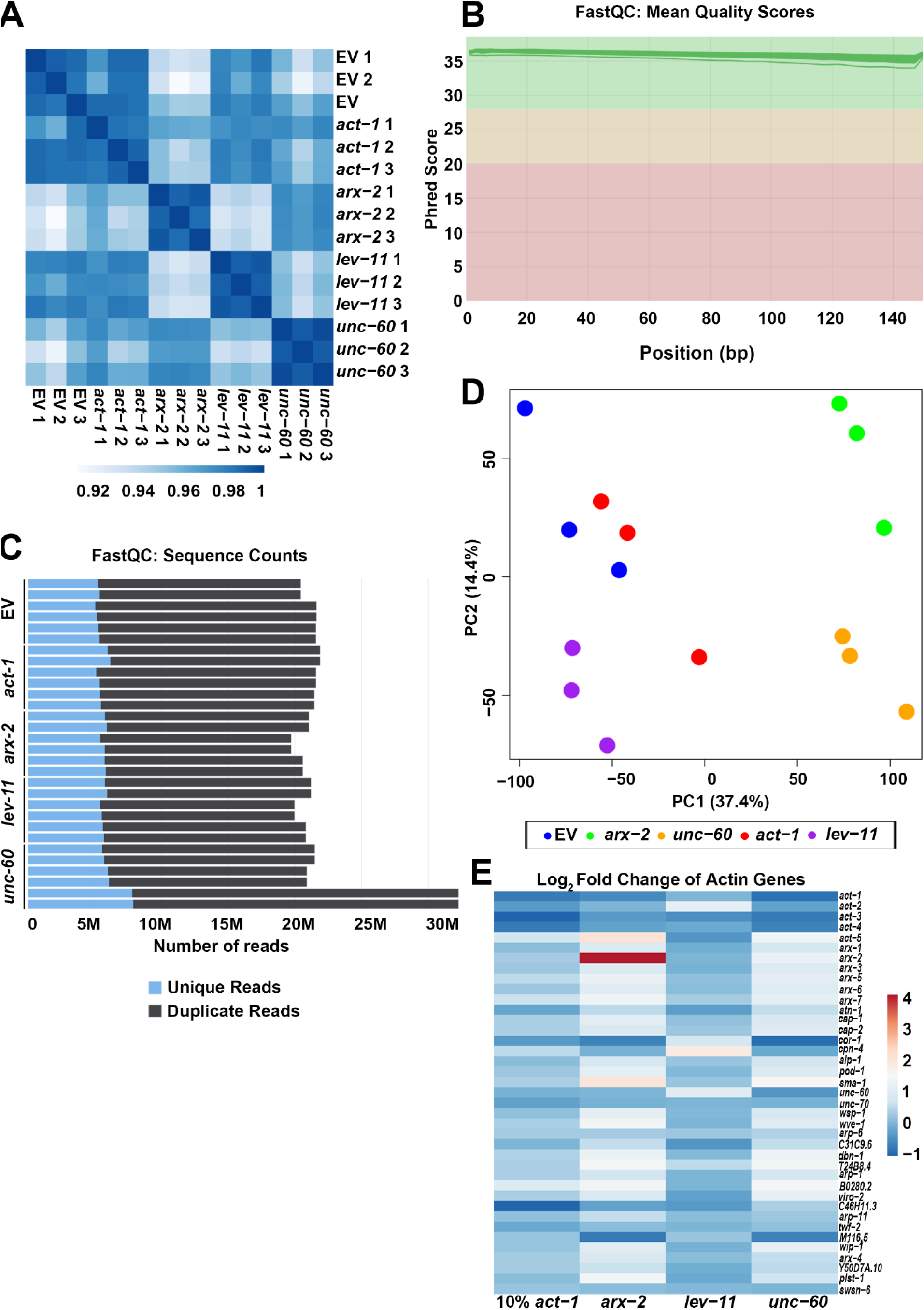
Quality control for RNA-seq libraries of C. elegans exposed to actin and ABP knockdown. **(A)** Spearman correlation plot of all RNA sequencing libraries. **(B)** Mean quality score (Phred score) of each RNA sequencing library. X-axis and Y-axis indicate the base pair position of each sequence and the Phred score, respectively. The graph was generated by MultiQC tool^189^. **(C)** The number of unique (blue) and duplicated (grey) reads from each pair-wise sequencing library (n=3). **(D)** PCA plots of actin and ABP knockdown in *C. elegans*. **(E)** Heatmap of log2(fold changes) for all genes annotated as cytoskeleton: Actin function in WormCat^190^. See **Table S9** for expression details of the genes used in the heatmap.

**Fig S2.**
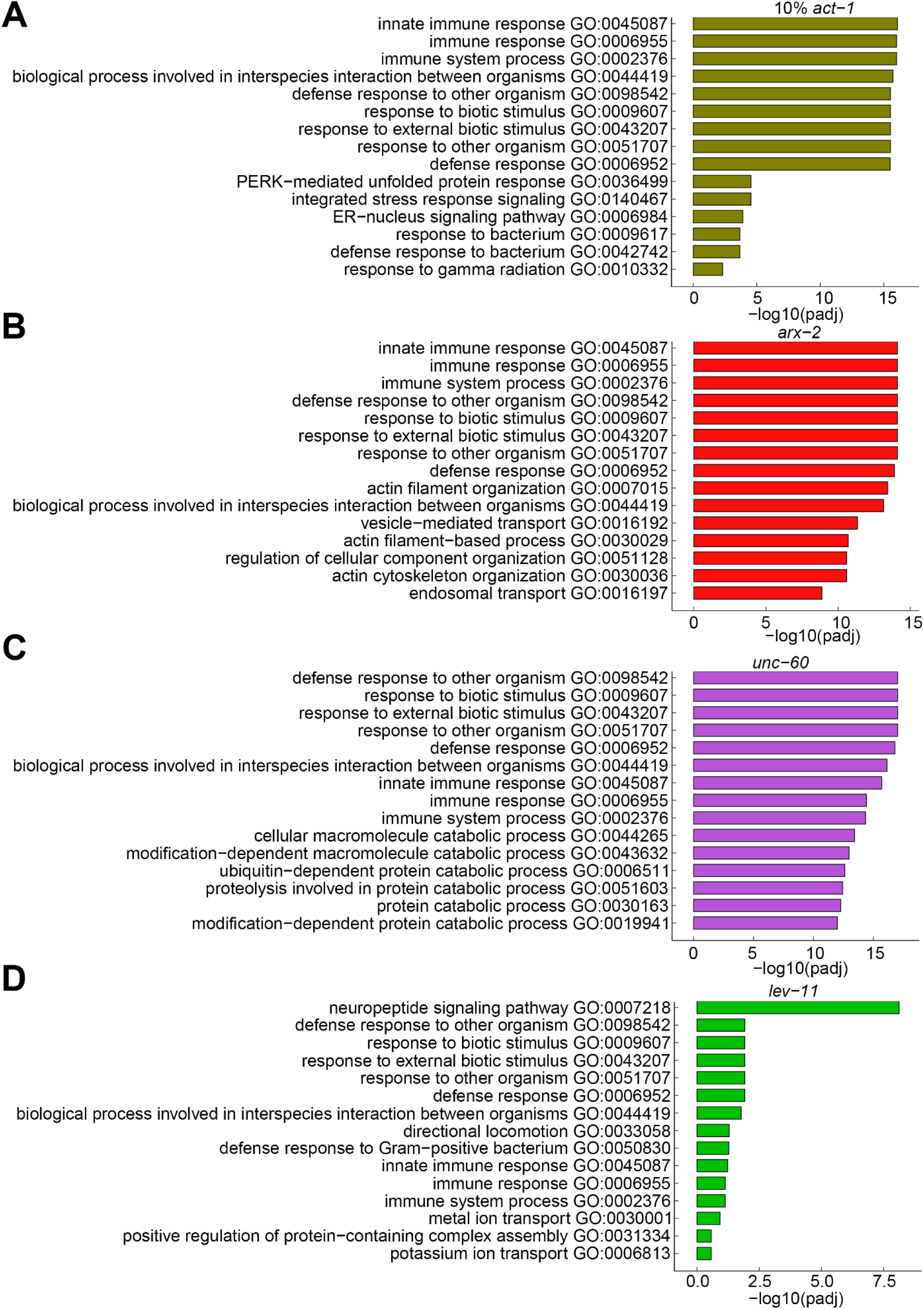
Gene ontology analysis of actin disruption. GO analysis of unique DEGs not shared with another condition for **(A)** *act-1,* **(B)** *arx-2***, (C**) *unc-60* and **(D)** *lev-11*. All DEGs were selected for adj p-val < 0.05, and the GOs were biological processes (BP) with q-value < 0.5. See **Table S5** for Go IDs, genes and statistics of the GO Analysis.

**Fig S3.**
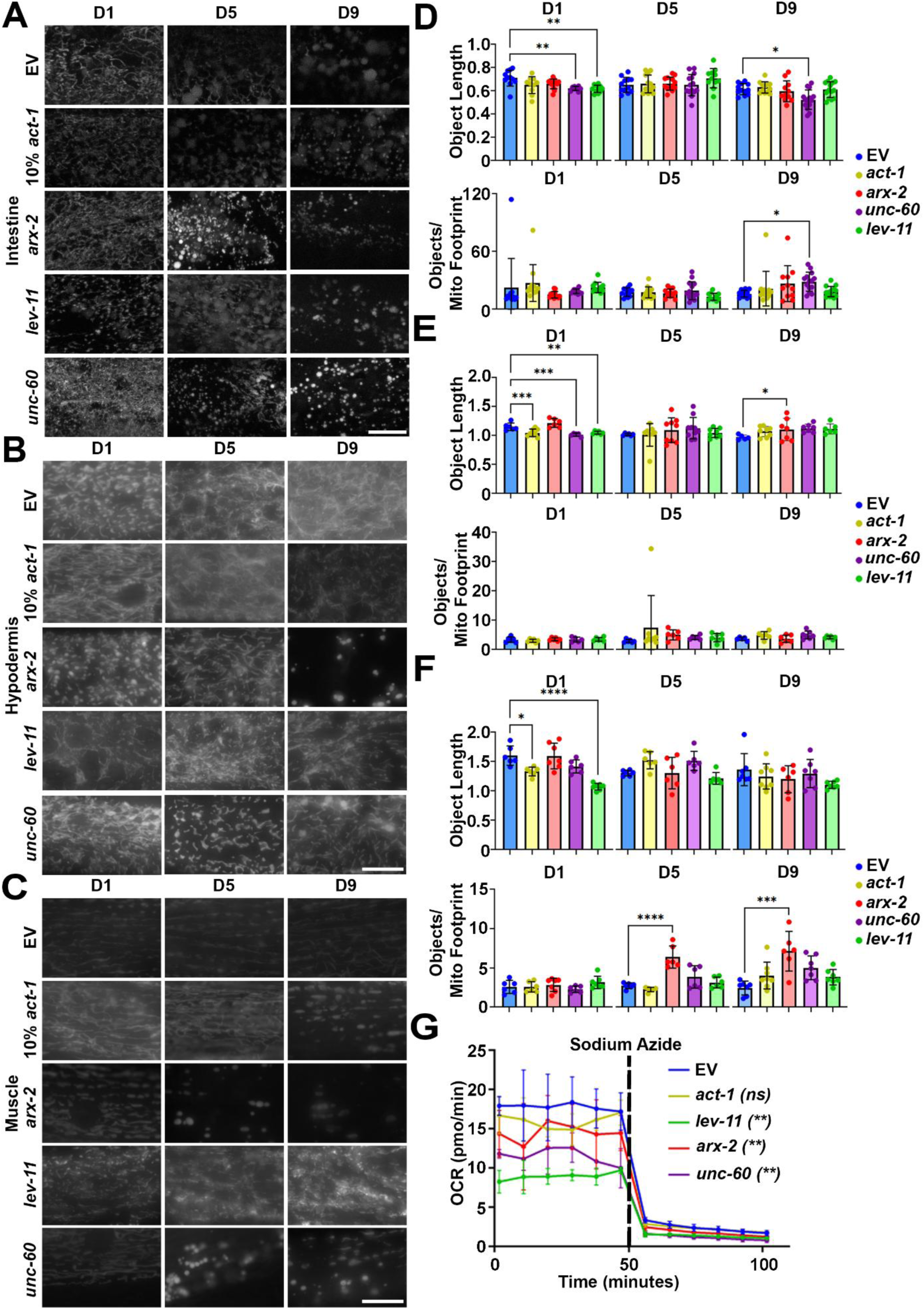
Actin disruption leads to mitochondrial dysfunction. Animals expressing *vha-6p::MLS::GFP* **(A)**, *col-19p::MLS::GFP* **(B)**, and *myo-3p::MLS::GFP* **(C)**, were grown on empty vector (EV), a 1:9 mix of *act-1*/EV (10% *act-1*), *arx-2*, *lev-11*, or *unc-60* RNAi from hatch and imaged during days 1, 5, and 9 of adulthood. **(D, E, F)** All quantification was performed using mitoMAPR. Object length and objects/mitochondrial footprint are shown here as example measurements, and all mitochondrial measurements measured by mitoMAPR are available in **Table S8**. In mitoMAPR-based quantification, the “objects/mitochondrial footprint” refers to the total number of objects detected as mitochondria and the area occupied by all the mitochondria within a defined region of interest**. (G)** Seahorse analysis of mitochondrial respiration/OCR (pmol/min) of day 1 wild type N2 animals grown on empty vector (EV, blue), 1:9 ratio of *act-1*:EV RNAi (yellow), *arx-2* (red), *lev-11* (green), or *unc-60* (purple) RNAi from hatch. 50 mM sodium azide was applied to measure non-mitochondrial respiration. Scale bar is 10 μm.

**Fig S4.**
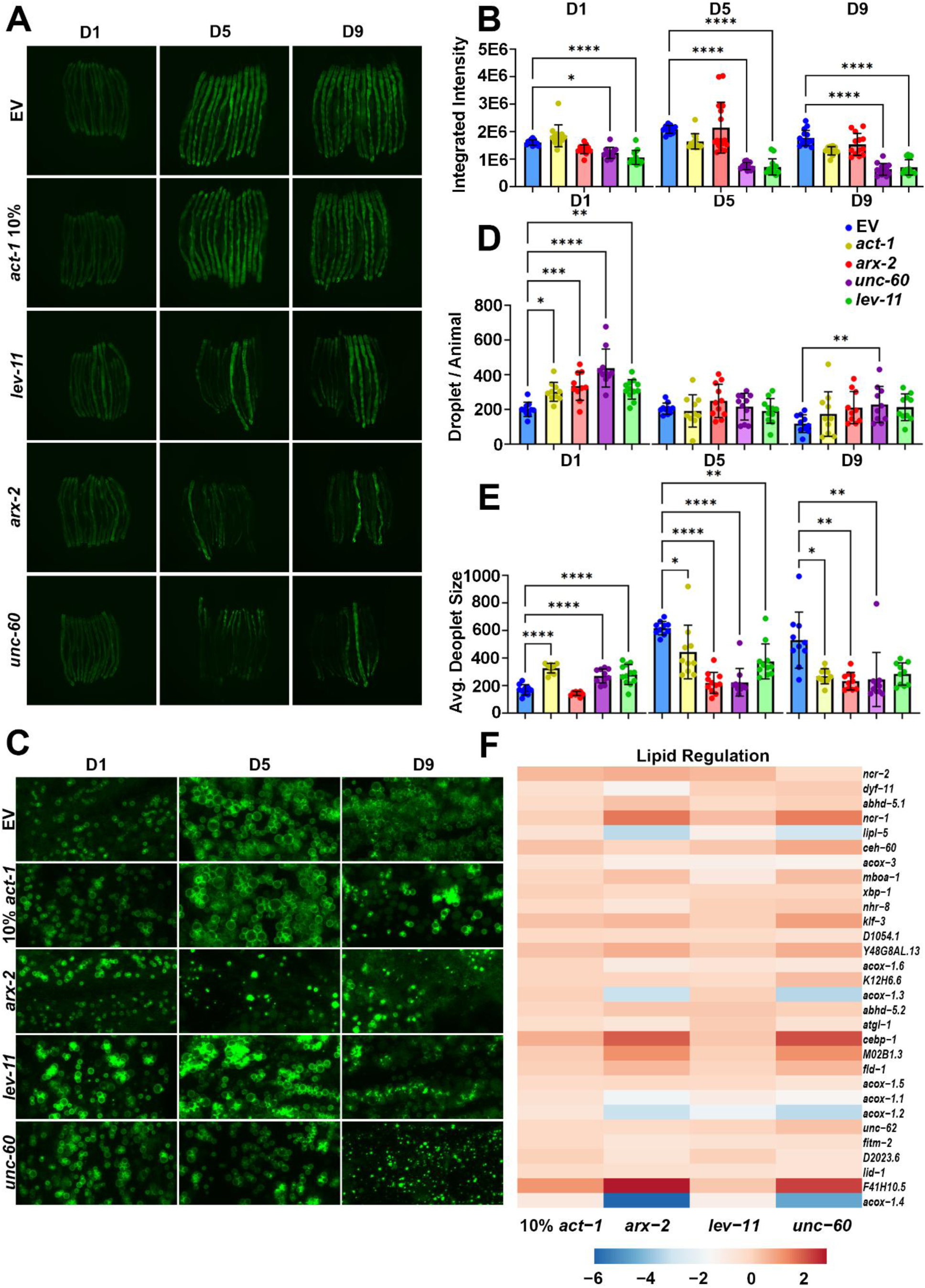
Actin disruption reduces lipid droplet size during aging. **(A)** Representative fluorescent stereomicroscope images of lipid droplets by visualization of DHS-3::GFP. Animals were grown on empty vector (EV), a 1:9 mix of *act-1*/EV (10% *act-1*), *arx-2*, *lev-11*, or *unc-60* RNAi from hatch. All animals were imaged on day 1, 5, and 9 of adulthood. **(B)** Quantification of GFP signal as measured by integrated intensity, based on 3 biological replicates of ≥ 12 animals per experimental condition. Two-way ANOVA was used for each time point. **(C)** Representative fluorescent confocal images of lipid droplets by visualization of DHS-3::GFP. Animals were grown on empty vector (EV), a 1:9 mix of *act-1*/EV (10% *act-1*), *arx-2*, *lev-11*, or *unc-60* RNAi from hatch. All animals were imaged on day 1, 5, and 9 of adulthood. **(D)** Quantification of the number of lipids per animal, based on 3 biological replicates of ≥ 4 animals per experimental condition. Two-way ANOVA was used for each time point. **(E)** Quantification of the average lipid size (in pixels) in each animal, based on 3 biological replicates of ≥ 4 animals per experimental condition. Two-way ANOVA was used for each time point. **(F)** Heat map of gene expression changes involved in lipid homeostasis (GO:0055088). See **Table S9** for expression details of the genes used in the heatmap.

**Fig S5.**
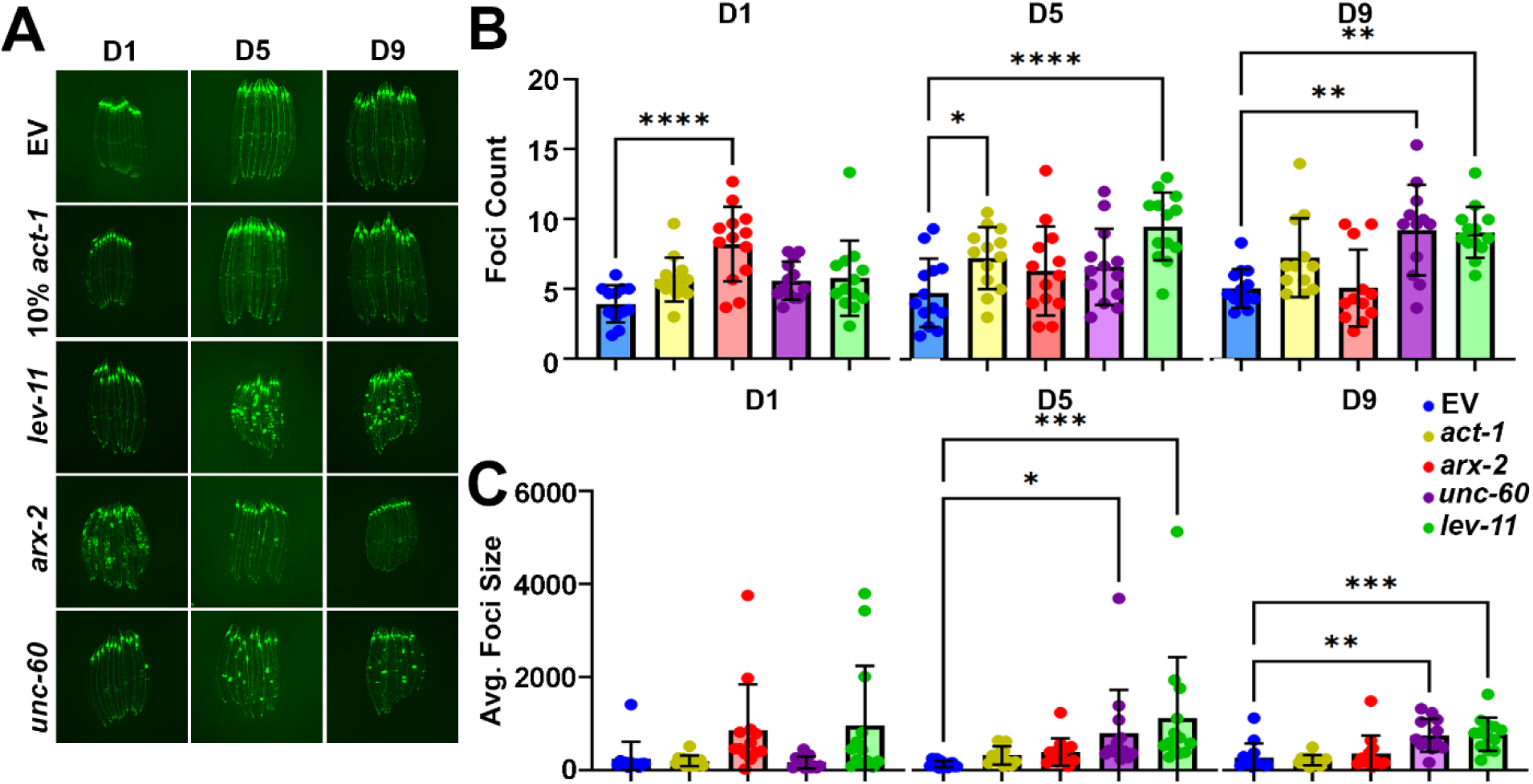
Actin disruption leads to dysregulation of proteostasis. **(A)** Representative stereomicroscope images of protein aggregation in animals expressing neuronal polyglutamine repeats (*rgef-1p::polyQ40::YFP*) at Day 1, Day 5 and Day 9 of adulthood. Animals were grown on empty vector (EV), a 1:9 mix of *act-1*/EV (10% *act-1*), *arx-2*, *lev-11*, or *unc-60* RNAi from hatch. Quantification of the **(B)** total number of aggregates and **(C)** average aggregate size was based on 3 biological replicates of ≥ 12 animals per experimental condition. Two-way ANOVA was used for each time point.

**Fig S6.**
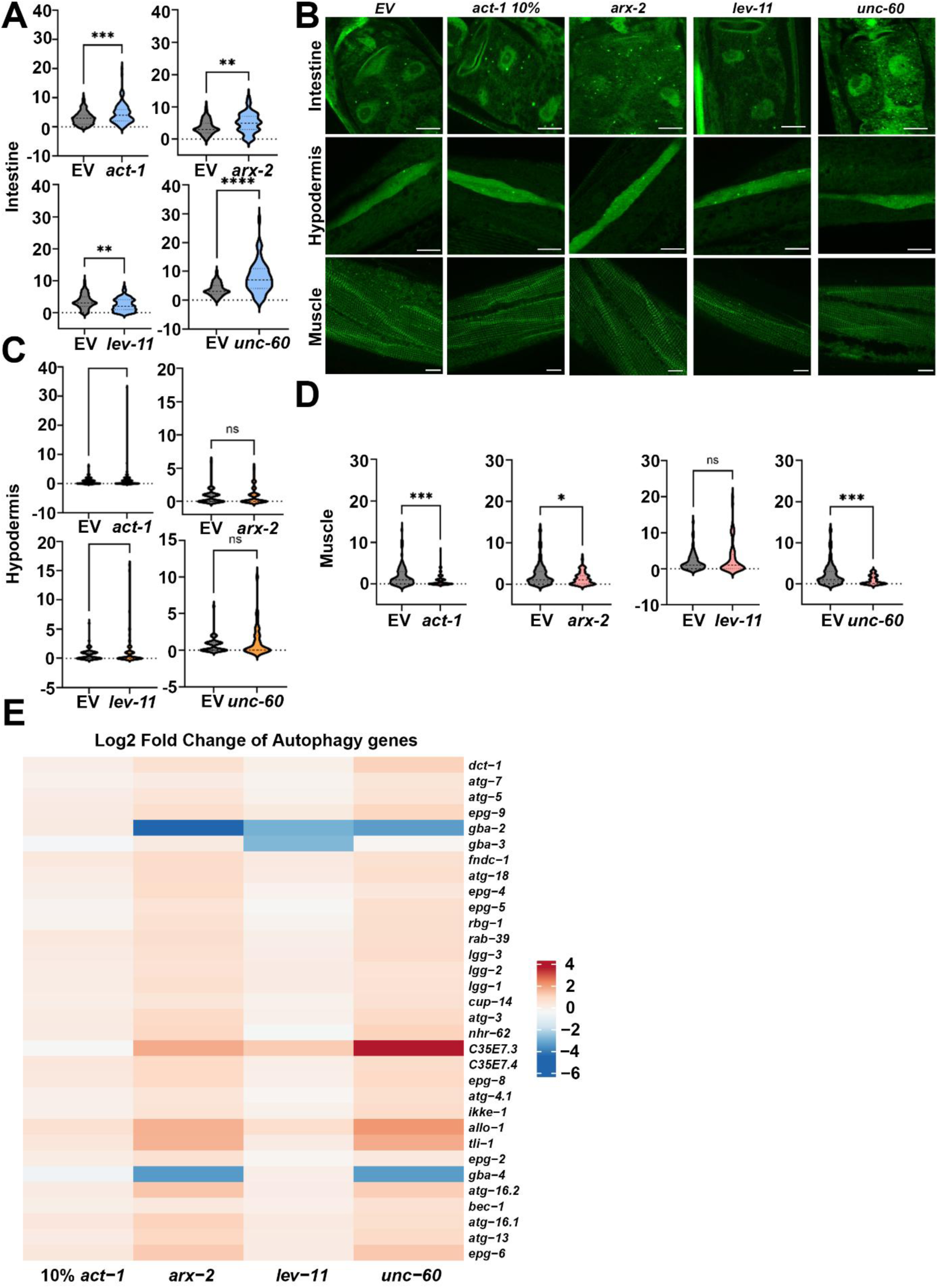
Actin disruption leads to tissue-specific dysregulation of autophagy. **(B)** Representative images of GFP::LGG-1 puncta in intestinal, hypodermal seam and muscle cells of control and RNAi-treated animals targeting *act-1, arx-2, unc-60, or lev-11* from hatch. Violin plots show quantification of GFP::LGG-1 puncta in **(A)** intestinal cells (CTRL, N = 63-107; *act-1*, N = 113; *arx-2*, N = 63; *unc-60*, N = 71; *lev-11*, N = 74 cells), **(C)** hypodermal seam cells (CTRL, N = 84–87; *act-1*, N = 87; *arx-2*, N = 51; *unc-60*, N = 45; *lev-11*, N = 70 cells) and **(D)** muscle cells (CTRL, N = 84–88; *act-1*, N = 88; *arx-2*, N = 51; *unc-60*, N = 37; *lev-11*, N = 57 cells. Data are pooled from three to five independent experiments. Statistical significance was determined by Welch’s two-tailed t-test (Graphpad). ns: P > 0.05, *P < 0.05, **P < 0.01, ***P < 0.001, ****P < 0.0001. Scale bars: 10 μm. **(E)** Heat map of the changes in expression (p<0.5) of genes involved in autophagy (directly annotated as “autophagy” in AmiGO2^188^). See **Table S9** for expression details of the genes used in the heatmap.

**Fig S7.**
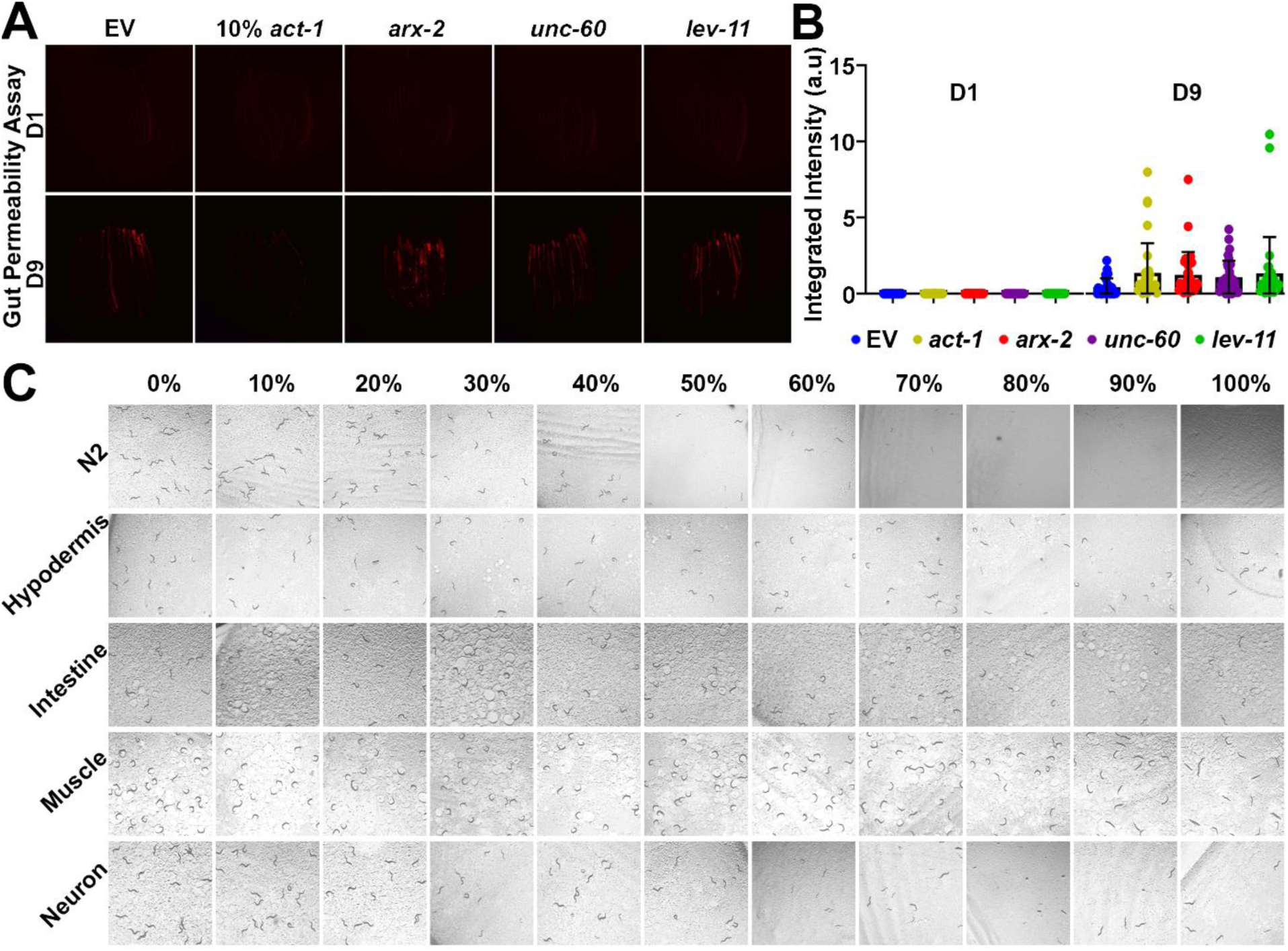
Actin disruption leads to tissue-specific dysregulation of gut barrier integrity. **(A)** Representative stereoscope images of gut colonization at Day 1 and Day 9 in animals grown with EV, 10% *act-1*, *arx-2*, *lev-11*, or *unc-60* RNAi mixed with 20% HT115 bacteria driving mCherry expression. **(B)** Quantification of gut colonization fluorescence signal. 10-15 worms were imaged for two technical replicates for each of 3 independent biological replicates (with ≥ 150 worms total per replicate per strain/condition), analyzed via two-way ANOVA test. **(C)** Representative stereoscopic images of *C. elegans* exposed to varying dilutions 0%-100% of actin RNAi from L1 stage of development. X-axis indicates dilution percentage with 0% being no actin RNAi and 100% meaning non-diluted RNAi. Y-axis represents the tissue where RNAi is functional with N2 being the wild-type control with whole-body functioning RNAi machinery.

**Fig S8.**
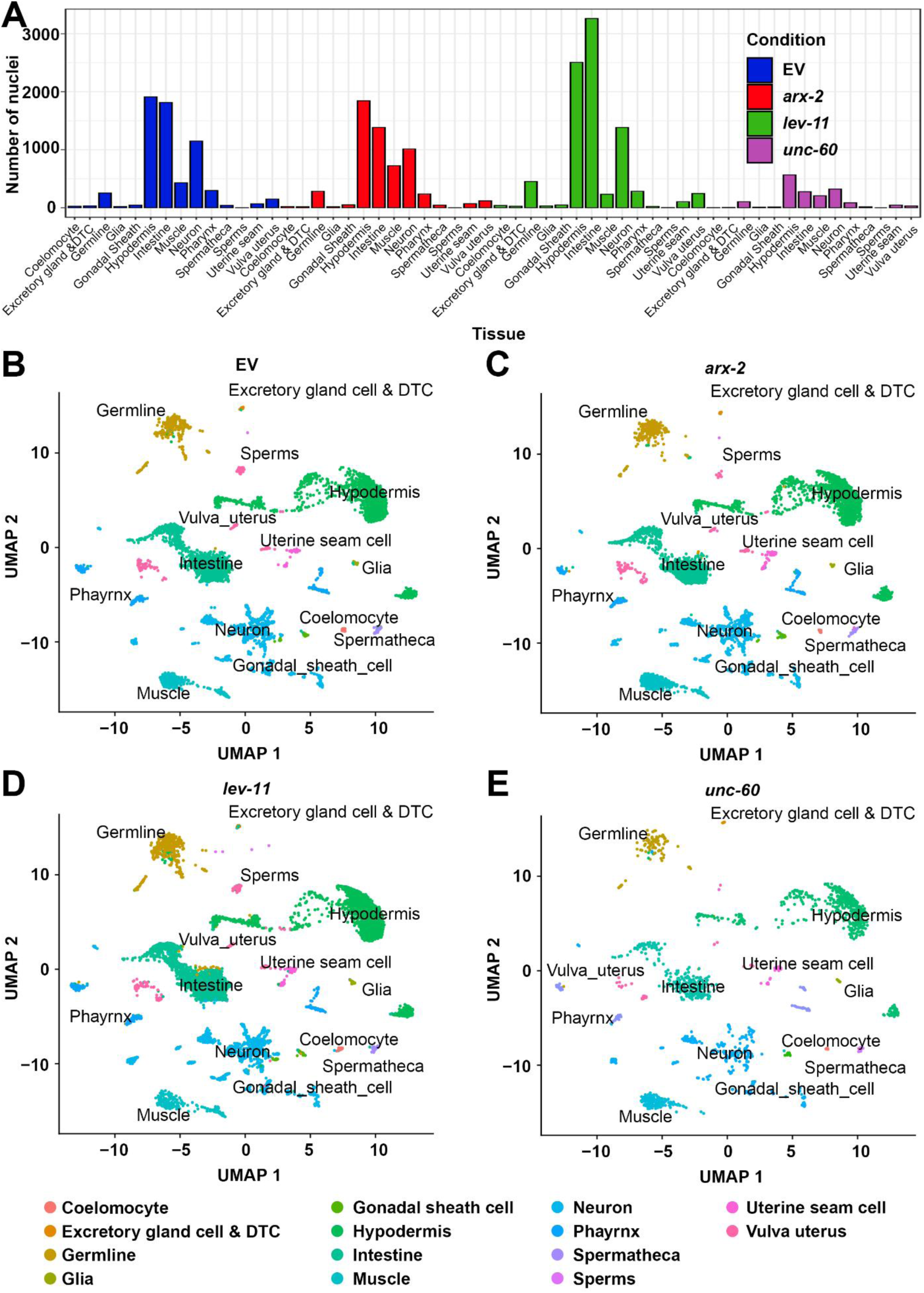
Annotation and clustering of single-nuclei RNA sequencing. **(A)** Summary plot of all sequenced annotated nuclei for each tissue and condition. **(B-E)** UMAP of the data segmented to corresponding cell types, as identified by cell-type biomarkers in D1 **(B)** control animals and animals with **(C)** *arx-2*, **(D)** *lev-11*, and **(E)** *unc-60* knockdown.

**Fig S9.**
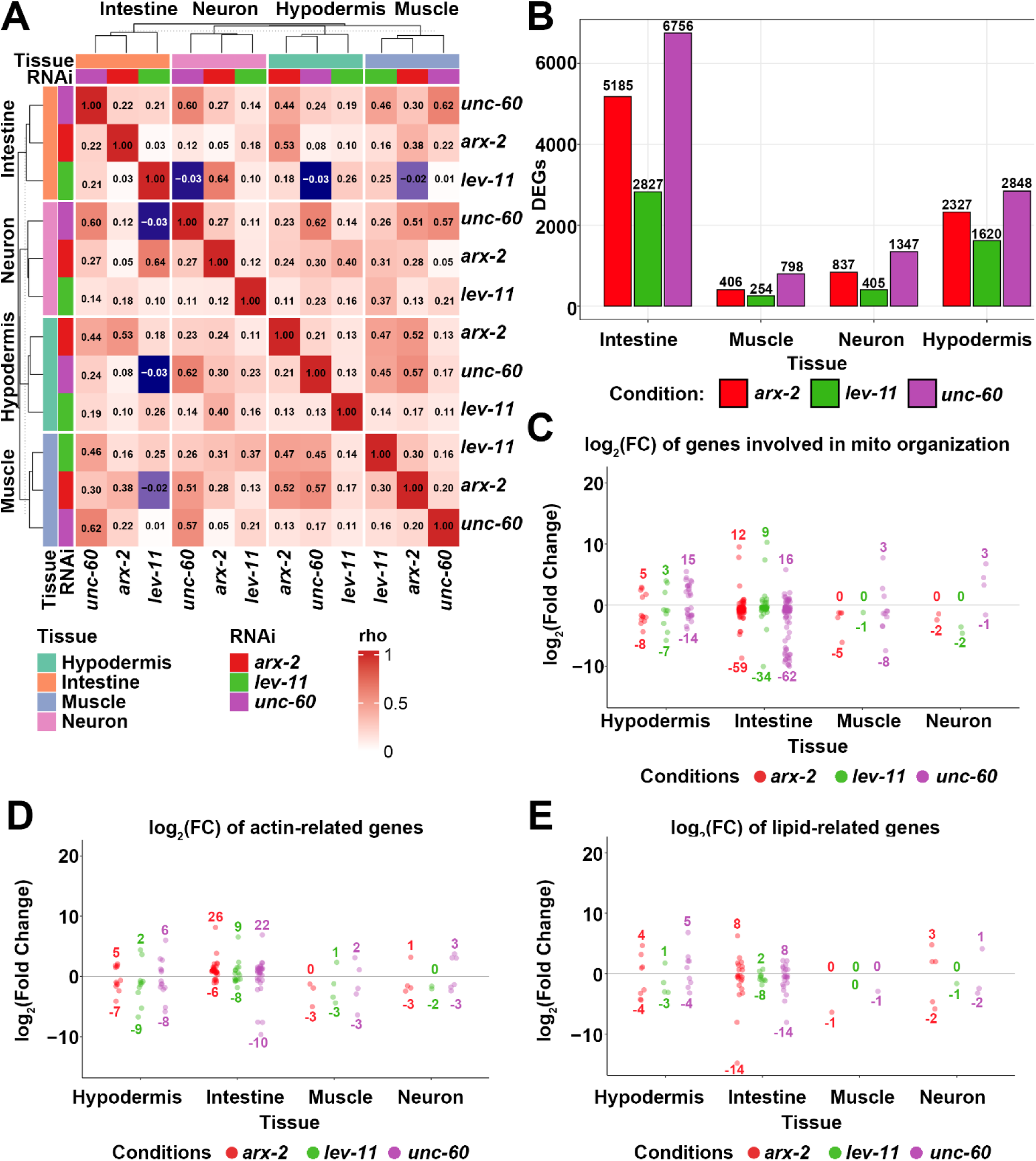
Deconvolution supplemental information. **(A)** Clustering of all samples based on the Spearman correlation between every tissue and condition. Clustering is performed within each tissue slice, and dendrograms illustrate the similarity structure. See **Table S10** for Spearman correlations between each sample. **(B)** Bar plot showing the number of DEGs for each condition and key selected tissues: Intestine, Muscle, Neuron and Hypodermis. (**C-E**) Dot plots showing significant DEGs involved in **(C)** mitochondrial organization (labeled as “involved in mitochondrion organization” in AmiGO2^188^), **(D)** actin maintenance (annotated as cytoskeleton: Actin function in WormCat) and **(E)** lipid regulation (GO:0055088) under ABP knockdown in 4 key tissues: Hypodermis, Intestine, Muscle and Neuron. See **Table S6** for DEGs in each dotplot.

**Fig S10.**
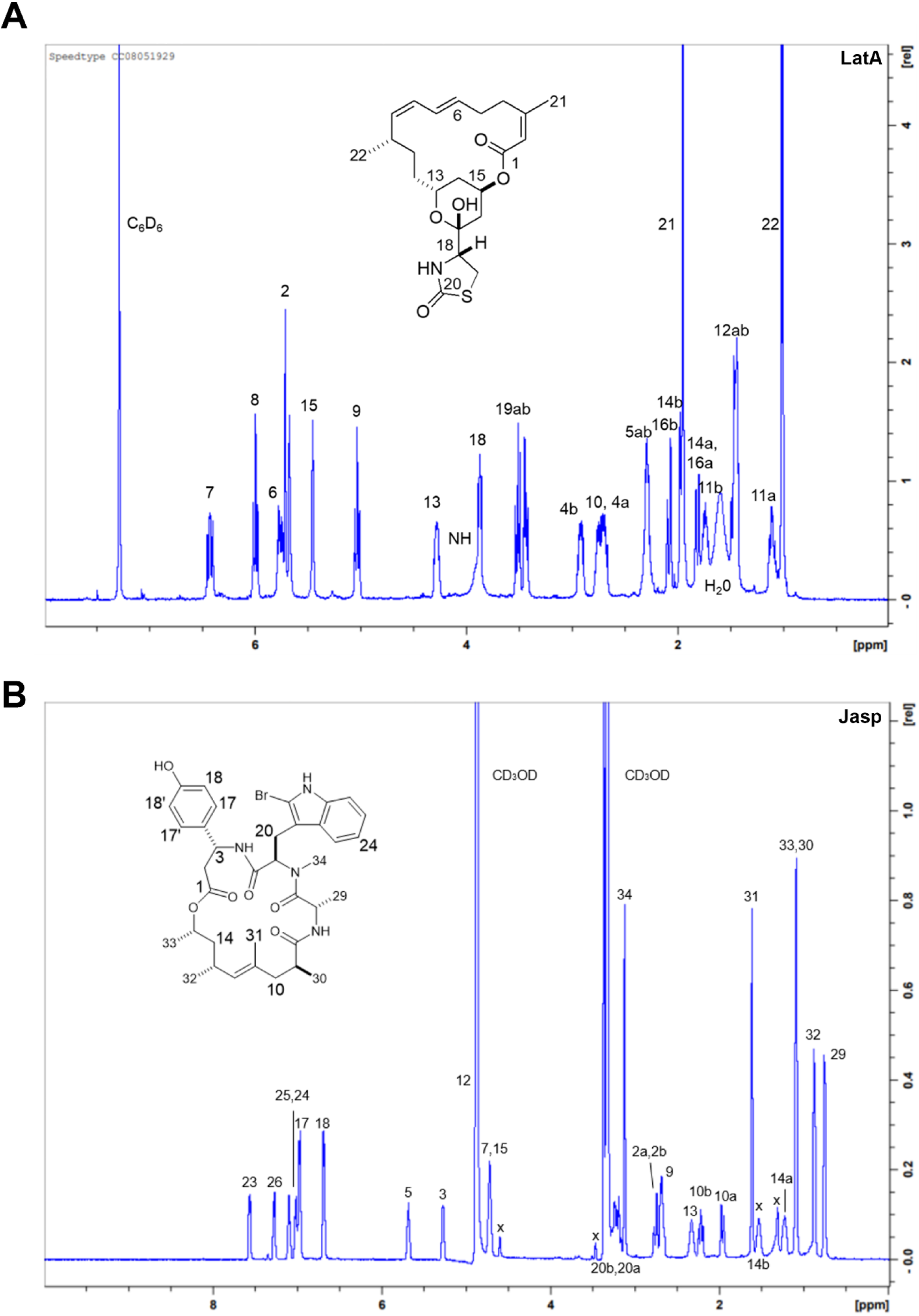
NMR analysis of LatA and Jasp. ^1^H NMR spectrum of **(A)** latrunculin A (LatA) in Benzene-*d6* at 500 MHz and **(B)** jasplakinolide (Jasp) in CD_3_OD at 500MHz

**Fig S11.**
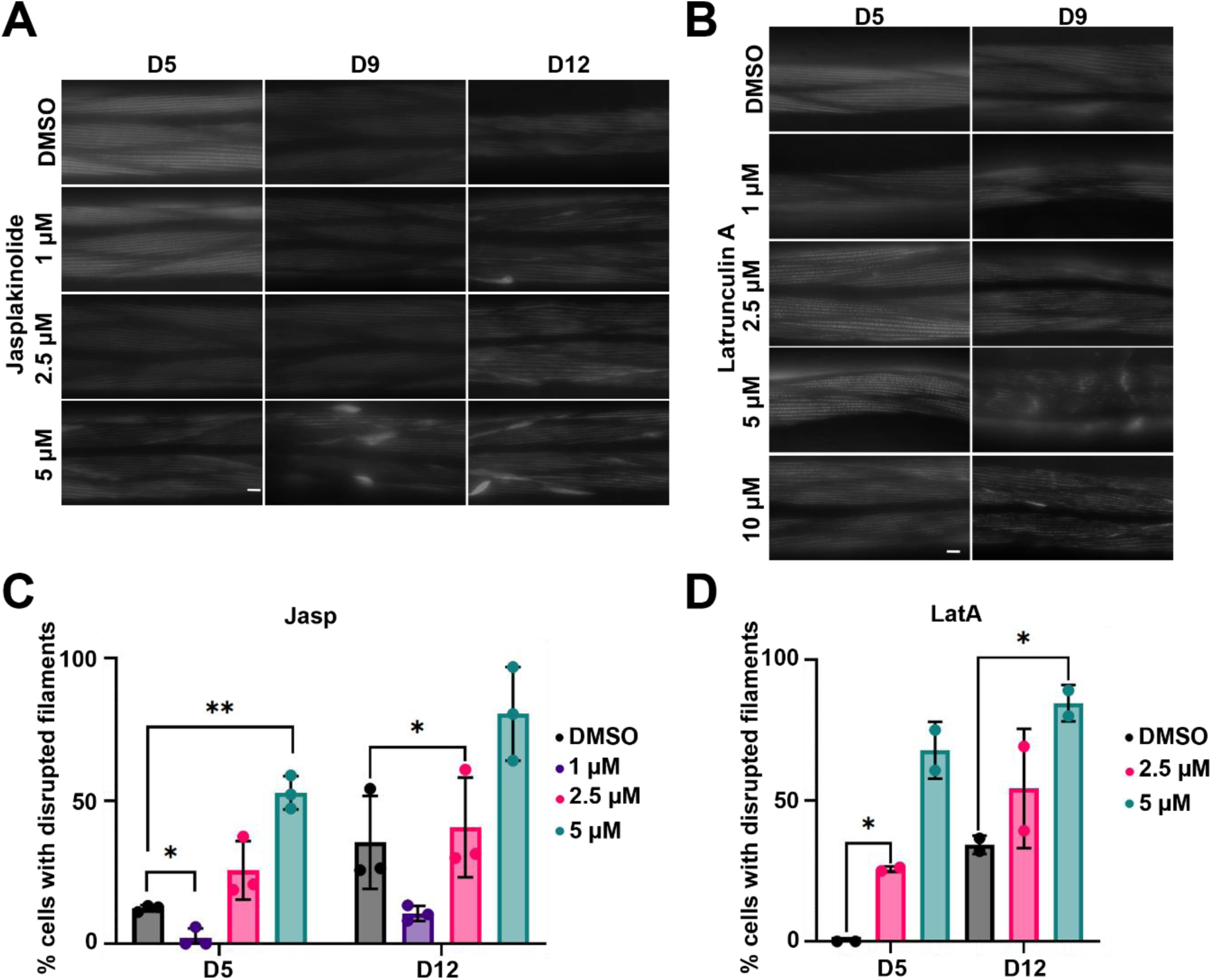
Small-molecule actin disruption effect on healthspan. **(A)** Representative fluorescent images of adult animals expressing LifeAct::mRuby in the muscle grown on empty vector (EV) and varying concentrations of Jasplakinolide (1µM, 2.5 µM, 5 µM) or DMSO control. **(B)** Representative fluorescent images of adult animals expressing LifeAct::mRuby in the muscle grown on empty vector (EV) and varying concentrations of Latrunculin A (1µM, 2.5 µM, 5 µM, 10 µM) or DMSO control. All muscle images were captured on day 5, and 9 of adulthood and on a Leica Thunder Imager. **(C-D)** Quantification of actin structure quality in animals expressing LifeAct::mRuby in the muscle grown on empty vector (EV) and varying concentrations of **(C)** Jasplakinolide and **(D)** Latrunculin A (1µM, 2.5 µM, 5 µM) or DMSO control. N=3 n>5.

**Fig S12.**
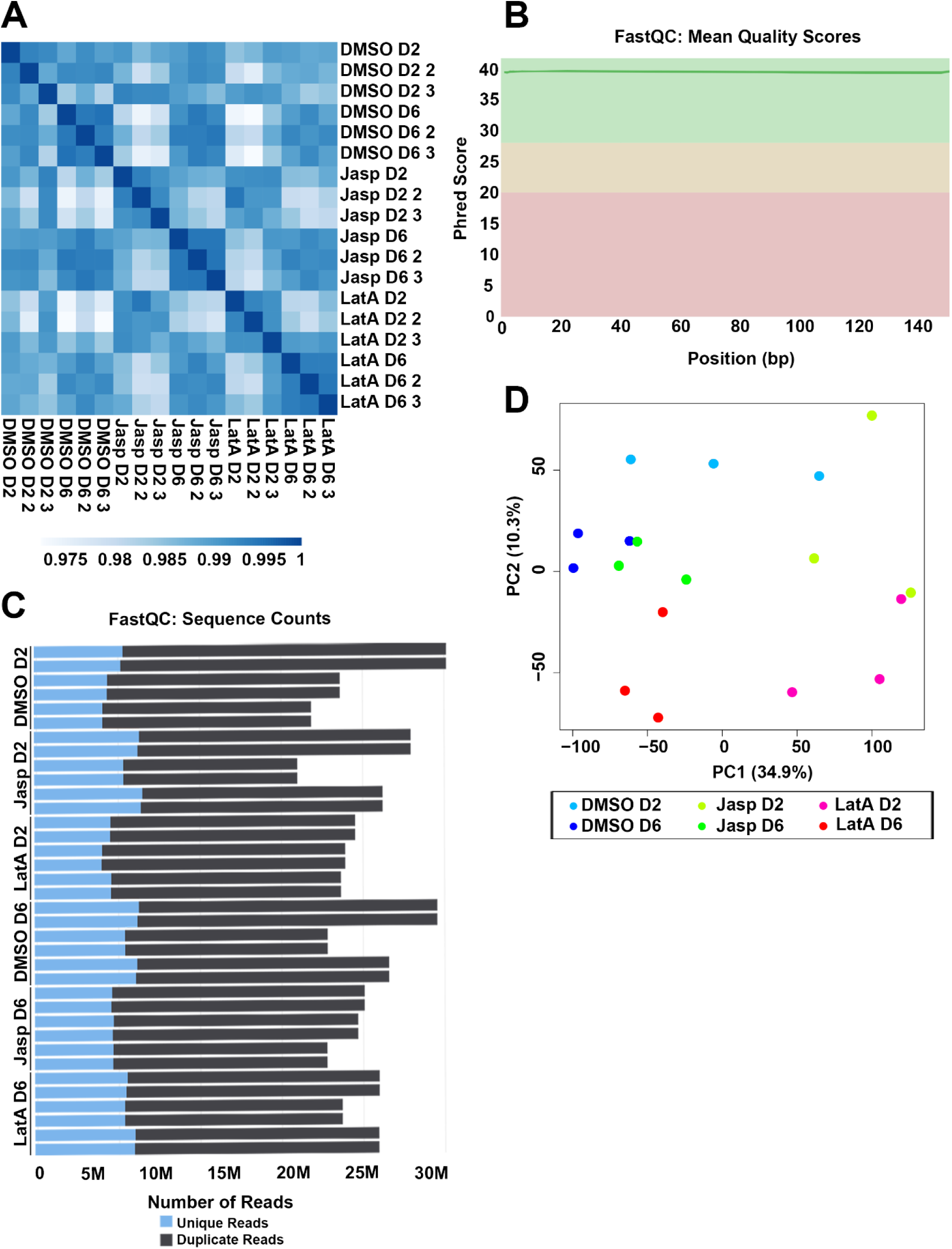
Quality control of LatA and Jasp RNA-seq data. **(A)** Spearman correlation plot of all RNA sequencing libraries. **(B)** Mean quality score (Phred score) of each sequencing library. X-axis and Y-axis indicate the base pair position of each sequence and the Phred score, respectively. The graph was generated by MultiQC tool^189^. **(C)** The number of unique (blue) and duplicated (grey) reads from each pair-wise sequencing library (n=3). **(D)** PCA plots of Jasplakinolide and Latrunculin A exposure in C. elegans.

**Fig S13.**
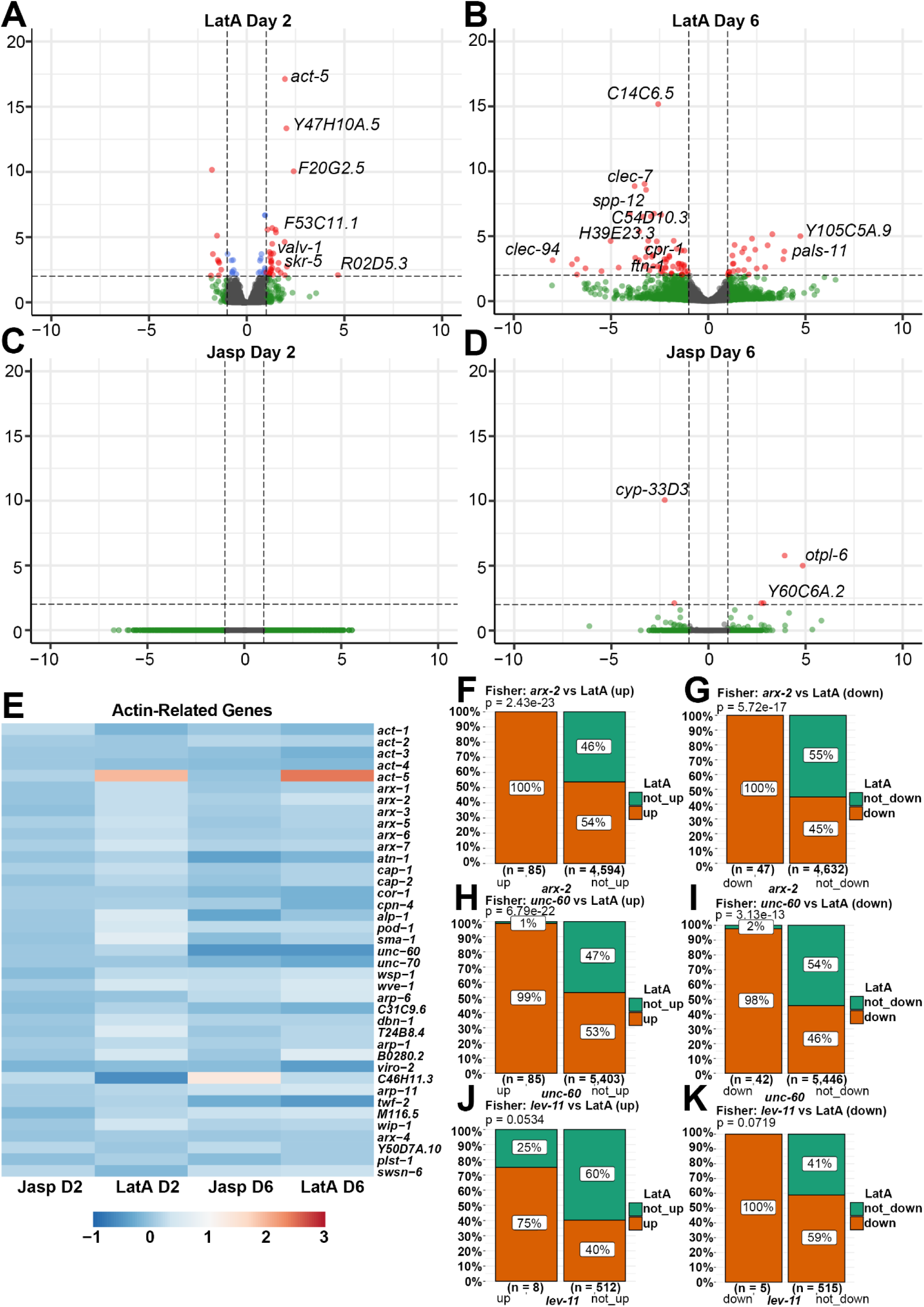
Actin-binding small molecules induces a mild transcriptional response. Volcano plots of genome-wide changes in gene expression upon drug exposure from Day 1 of adulthood on LatA until **(A)** Day 2 and **(B)** Day 6, and Jasp until **(C)** Day 2 and **(D)** Day 6 of adulthood. Red dots indicate significantly differentially expressed genes with p-value ≤ 0.01 and log_2_FC>|2|. Blue dots indicate significantly differentially expressed genes with p-value ≤ 0.01 and log_2_FC<|2|. Green dots indicate significantly differentially expressed genes with p-value ≥0.01. See **Table S11** for a list of differentially expressed genes and expression values. **(E)** Heat map of differentially expressed genes annotated as “cytoskeleton: Actin function” in WormCat^190^. Warmer colors indicate increased expression, and cooler colors indicate decreased expression. See **Table S9** for expression details of the genes used in the heatmap. **(F-K)** Bar plots to visualize the overlap of up- and down-regulated differentially expressed genes (DEGs) between animals treated with **(F)(G)** *arx-2* and LatA, **(H)(I)** *unc-60* and LatA, and **(J)(K)** *lev-11* and LatA, using ggbarstats function in R. Statistical significance of the overlap was assessed using Fisher’s exact test.

