## Supplementary material for "Form and function of actin impacts actin health and aging": Source Code

**A.) .IJM code for hypodermal actin quantification**

run("Clear Results"); // clear the results table of any previous measurements

setBatchMode(true);

inputDirectory = getDirectory("Choose a Directory of Images");

fileList = getFileList(inputDirectory);

for (i = 0; i < fileList.length; i++)

{

processImage(fileList[i]);

}

setBatchMode(false); // Now disable BatchMode since we are finished

updateResults(); // Update the results table so it shows the filenames

// Show a dialog to allow user to save the results file

outputFile = File.openDialog("Save results file");

// Save the results data

saveAs("results",outputFile);

function processImage(imageFile)

{

// Store the number of results before executing the commands,

// so we can add the filename just to the new results

prevNumResults = nResults;

open(imageFile);

// Get the filename from the title of the image that's open for adding to the results table

// We do this instead of using the imageFile parameter so that the

// directory path is not included on the table

filename = getTitle();

run("8-bit");

setAutoThreshold("Default dark no-reset");

run("Convert to Mask");

run("Set Measurements...", "area mean min area_fraction redirect=None decimal=3");

run("Measure");

for (row = prevNumResults; row < nResults; row++)

{

setResult("Filename", row, filename);

}

close("*"); // Closes all images

}

**B.) .IJM code for lipid droplet quantification**

// Close existing Results table if open

if (isOpen("Results")) {

selectWindow("Results");

run("Close");

}

// Prepare a dummy measurement to initialize Results table

run("Set Measurements...", "area mean integrated redirect=None decimal=3");

run("Measure");

setResult("Label", 0, "Init");

updateResults();

run("Clear Results");

for (img = 0; img < nImages; img++) {

selectImage(img + 1);

title = getTitle();

// Switch to channel 2

Stack.setChannel(2);

run("8-bit");

run("Enhance Contrast", "saturated=0.35");

// Prompt for ROI input

roiManager("reset");

waitForUser("Image: " + title + " (Channel 2)\nDraw ROIs around each worm and add them to the ROI Manager.\nClick OK to continue.");

nWorms = roiManager("count");

for (i = 0; i < nWorms; i++) {

roiManager("Select", i);

run("Measure");

// Ensure the Results table is selected before editing

selectWindow("Results");

currentRow = nResults - 1;

label = "Worm_" + (i+1) + "_" + title;

setResult("Label", currentRow, label);

updateResults();

}

}

// Ask user where to save the file

saveDir = getDirectory("Choose a folder to save your CSV");

filePath = saveDir + "IntegratedIntensity_Summary_sqstR3.csv";

// Save the results

selectWindow("Results");

saveAs("Results", filePath);

**C.).IJM code for gut assay quantification**

// Close existing Results table if open

if (isOpen("Results")) {

selectWindow("Results");

run("Close");

}

// Prepare a dummy measurement to initialize Results table

run("Set Measurements...", "area mean integrated limit redirect=None decimal=3");

run("Measure");

setResult("Label", 0, "Init");

updateResults();

run("Clear Results");

for (img = 0; img < nImages; img++) {

selectImage(img + 1);

title = getTitle();

// Switch to channel 2

Stack.setChannel(2);

run("8-bit");

// run("Enhance Contrast", "saturated=0.35");

// Prompt for ROI input

roiManager("reset");

waitForUser("Image: " + title + " (Channel 2)\nDraw ROIs around each worm and add them to the ROI Manager.\nClick OK to continue.");

nWorms = roiManager("count");

for (i = 0; i < nWorms; i++) {

roiManager("Select", i);

Stack.setChannel(2);

setThreshold(25, 255); // Only include pixels above 25

setOption("BlackBackground", true);

run("Measure");

// Ensure the Results table is selected before editing

selectWindow("Results");

currentRow = nResults - 1;

label = "Worm_" + (i+1) + "_" + title;

setResult("Label", currentRow, label);

updateResults();

}

}

// Ask user where to save the file

saveDir = getDirectory("Choose a folder to save your CSV");

filePath = saveDir + "dayx_replicatex _IntegratedIntensity_Summary_gut_assay.csv";

// Save the results

selectWindow("Results");

saveAs("Results", filePath);

**D.) .IJM code for lipid droplet and neuronal puncta quantification**

if (isOpen("Summary")) { selectWindow("Summary"); run("Close"); }

for (img = 0; img < nImages; img++) {

selectImage(img + 1);

title = getTitle();

// Switch to channel 3

Stack.setChannel(3);

// Prompt user to draw worm ROIs

roiManager("reset");

waitForUser("Image: " + title + " (Channel 3) \nDraw ROIs around each worm and add them to the ROI Manager.\nClick OK to continue.");

nWorms = roiManager("count");

for (i = 0; i < nWorms; i++) {

// Select ROI and duplicate region

roiManager("Select", i);

newTitle = "Worm_" + (i+1) + "_" + title;

run("Duplicate...", "title=" + newTitle);

actualTitle = getTitle();

selectWindow(actualTitle);

// Reapply ROI before clearing

roiManager("Select", i);

run("Clear Outside");

run("8-bit");

run("Subtract Background...", "rolling=30");

run("Auto Threshold", "method=MaxEntropy");

setOption("BlackBackground", true);

run("Convert to Mask");

run("Erode");

run("Dilate");

//waitForUser("Review thresholding for " + newTitle + ". Zoom/pan if needed, then click OK to continue.");

// Analyze particles into summary table with custom name

run("Set Measurements...", "area mean integrated redirect=None decimal=3");

run("Analyze Particles...", "size=0-Infinity circularity=0.0-1.00 summarize");

// Rename the Slice label in Summary table

sliceName = "Worm_" + (i+1) + "_" + title;

Table.set("Slice", Table.size - 1, sliceName);

close();

}

}

// Save the summary table to a folder of your choice

saveDir = getDirectory("Choose a folder to save your CSV");

filePath = saveDir + "Counts_SummaryTable.csv";

selectWindow("Summary");

saveAs("Results", filePath);
